# Interallelic cis-regulatory dominance promotes robustness and evolutionary innovation

**DOI:** 10.64898/2026.03.17.712157

**Authors:** Noa O Borst, Timothy Fuqua, Fabian Ruperti, Justin Crocker

## Abstract

Dominance is a central principle of genetics, yet the mechanistic basis and the evolutionary consequences of dominance arising from cis-regulatory variation remain poorly understood. We examined the evolutionary trajectories of a pleiotropic developmental enhancer in *Drosophila*. A genotype–phenotype map between *D. melanogaster* and *D. simulans* enhancer sequences reveals extensive epistasis, and many homozygous evolutionary paths reduce transcriptional output. In heterozygotes, however, regulatory dominance masks variants that reduce gene expression, potentially relaxing evolutionary constraints. Using allele-specific reporters and imaging, we show that this dominance arises from interallelic interactions (also known as transvection) reinforced by transcriptional hubs. Importantly, this enhancer dominance is cell-type specific, raising the possibility that it conceals deleterious effects in essential tissues while revealing novel, ectopic activity in others. Interallelic regulatory hubs may therefore expand the range of mutational paths available to diploid genomes while preserving essential transcriptional output.

## Quote

> “There are those who flatly denied the possibility of such a thing; (…) even decreed that it was heresy to believe that witches are conveyed from one place to another in the manner they allege.”

> — Henri Boguet, *An Examen of Witches (1602; English transl. 1929)*, Chapter XIV: *Of the Transvection of Witches to the Sabbat*

## Main

Dominance is a fundamental principle of genetics describing how the non-additive contributions of alleles shape the mapping from genotype to phenotype^1^. In protein-coding genes, dominance is often explained by dosage compensation or functional redundancy^1^, which can be beneficial in cases such as hybrid vigor^2^. While dominance has been extensively studied for protein-coding variation, the mechanistic basis of dominance arising from noncoding cis-regulatory sequences remains poorly understood^3^. Developmental enhancers, which integrate multiple transcription factor (TF) inputs to drive precise spatiotemporal gene expression^4^, provide a tractable system to explore this problem. Their activity depends on cooperative TF binding and the formation of transcriptional hubs^5^, which are transient high local concentrations of regulatory factors near enhancers^6^. Enhancer variants are considered a major source of morphological variation, primarily because they are thought to have fewer pleiotropic effects than protein-coding variants^7,8^.

Paradoxically, developmental enhancers appear both fragile and robust. Saturation mutagenesis studies across Metazoa indicate that many individual nucleotides are essential for correct enhancer activity^9–13^, implying a high degree of mutational fragility. Yet, across evolution, enhancers with extensive sequence divergence frequently maintain conserved expression patterns^14–16^, implying that regulatory output can be preserved despite sequence turnover. This fragility–robustness paradox^17^ raises a central question: how can enhancers accumulate mutations and acquire new functions without compromising developmental programs?

One possible explanation lies in the diploid nature of many eukaryotic genomes. Since new mutations arise in single copies, natural selection initially acts on heterozygous genotypes^18,19^. In this context, each cell carries two homologous enhancer alleles that can, in some cases, interact. In *Drosophila*, homologous chromosomes are somatically paired, enabling direct enhancer–promoter communication between alleles—a phenomenon historically termed transvection^20^. Similar interallelic and interchromosomal interactions have been documented across diverse organisms^21–24^, including mammals^25–27^, where enhancers and promoters can transiently share transcriptional hubs^6,28–30^. Such interallelic crosstalk within shared transcriptional environments could allow a functional allele to buffer transcriptional output from a mutant counterpart, thereby generating dominance at the level of regulatory architecture rather than protein function.

## Results

To test our hypothesis of interallelic dominance, we focused on the 292 base pair (bp) *E3N* enhancer of the *D. melanogaster shavenbaby* (*svb*; also known as *ovo*) gene, which drives segmented embryonic expression in ventral ectodermal cells and is required for cell differentiation in the larval ventral denticle belts^31,32^. We selected *E3N* because its function is largely conserved across *Drosophila* species despite substantial sequence divergence^31^. At the same time, saturation mutagenesis experiments have shown that random mutations in *E3N* can result in significant and pleiotropic changes in embryonic spatiotemporal gene expression patterns^12^, providing an ideal system to explore how a seemingly fragile enhancer could retain regulatory function throughout its evolutionary history.

### Functional and sequence divergence of the E3N enhancer within and between Drosophila species

To explore natural phenotypic variation in *E3N,* we assessed sequence variation in 1,121 wild-derived *D. melanogaster* genomes^33^ (**Fig. 1a,b, File S1**). Only seven polymorphism*s* occurred at frequencies ≥1%, suggesting limited sequence variation in the enhancer (**Fig. 1b,c**). *E3N* sequences containing these single nucleotide polymorphisms (SNPs) were cloned in a *lacZ* reporter plasmid and integrated into the *D. melanogaster* genome (see **Methods**). We tested the effect of each of these SNPs on the activity of the enhancer and found that, unlike random mutations—which typically decrease *E3N* activity^12,34^,—naturally occurring SNPs produced expression patterns comparable to the wild-type enhancer (from here on referred to as *E3N-lacZ,* **Fig. 1d-o, S2a**). To exclude buffering by native flanking sequences, we introduced a subset of variants into an extended 392 bp enhancer construct and observed comparable effects on enhancer activity (**Fig. S1**). Together, these results suggest that *E3N* is under selective pressure to maintain its regulatory activity.

**Figure 1.**
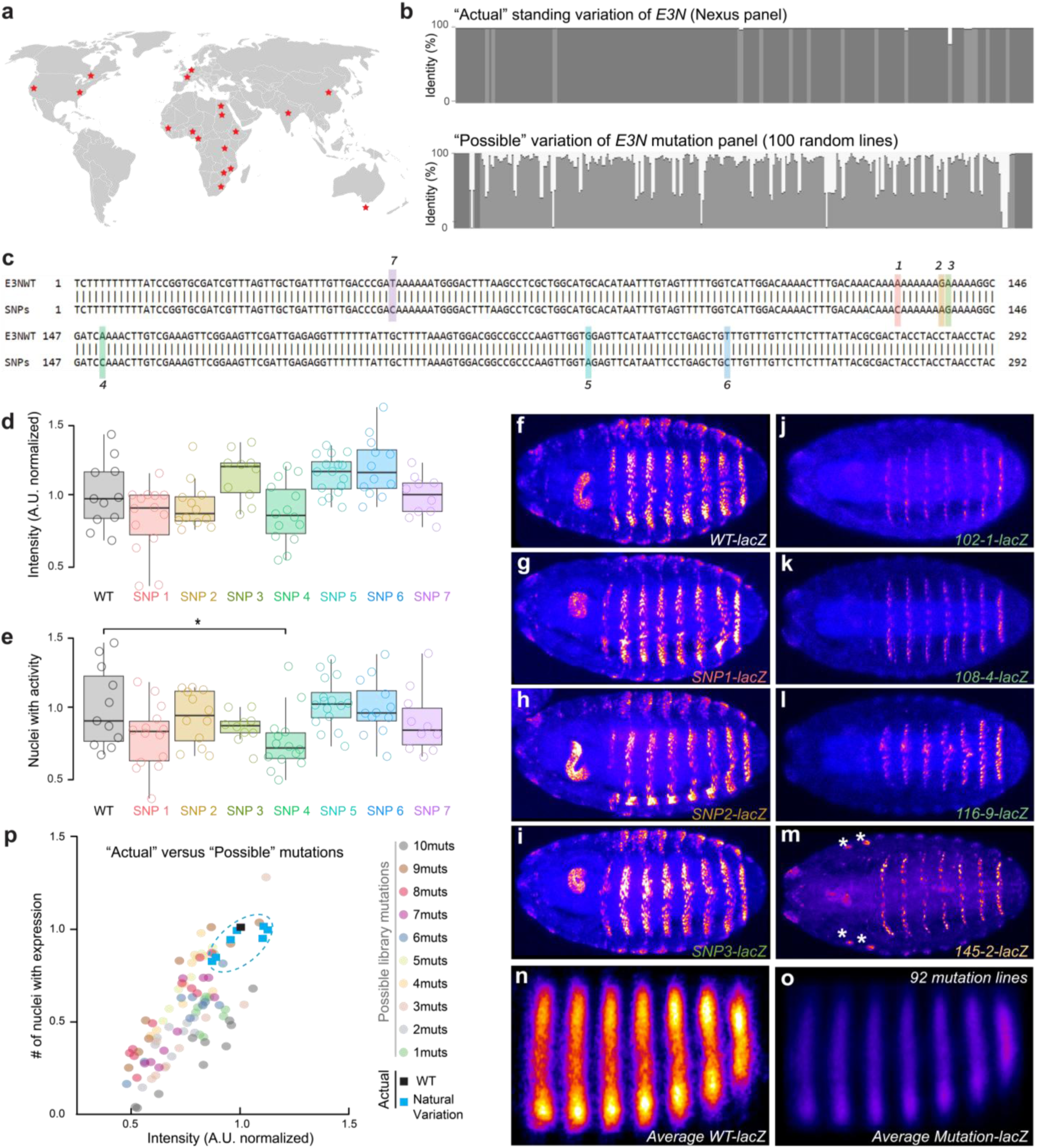
SNPs from global standing variation in *E3N* have little effect on the enhancer activity compared to random synthetic variants. (**a**) An overview of sample locations of the 1,121 Drosophila melanogaster genomes for the Drosophila Genome Nexus dataset^33^. **(b)** A comparison of the global standing variation in *E3N* in D. melanogaster (“the actual”) with the variation of a mutant *E3N* panel of 91 variants ranging from 1-10 mutations, taken from a random mutagenesis library of the enhancer^12,34^ (“the possible”). **(c)** Variations from the consensus sequence (E3NWT) occurring at a total rate of ≥ 1% in the Nexus dataset were considered hits for standing variation, and are labelled as SNP1-7. **(d)** Intensity of the gene expression patterns of the enhancer variants containing each of seven SNPs upon being inserted in a pLacZ::attB vector and integrated at the *attP2* landing site of the *D. melanogaster* genome. ANOVA with Dunnett’s post hoc test was performed between WT and all the individual SNPs, where no variant significantly different in its reporter gene intensity from WT. **(e)** Boxplot of the number of nuclei with activity of the enhancer variants containing each of seven SNPs. ANOVA with Dunnett’s post hoc test was performed between WT and all the individual SNPs, where SNP 4 proved significantly different from the wildtype expression pattern (p = 0.0259). (**f-m**) A selection of the different stripe patterns shown in (e), where (f) is the wildtype pattern, (g - i) are three SNPs from the 7 SNPs shown in d, (j - l) are examples of mutant embryos from the 91 mutant lines, and (m) is a mutant with ectopic expression in the wing and haltere discs. (**n-o**) An elastic registration of the embryos with WT *E3N* patterns (m) and all the embryos from the 91 mutation lines (o), showing the negative effect of the “average” mutation on the expression of the *lacZ* reporter gene, which serves as a read-out of the activity of the *E3N* enhancer. (**p**) A comparison of the effect of the natural variation and the 91 mutant variants on the average intensity of, and the number of nuclei with *lacZ* reporter expression.

We next examined divergence in *E3N* sequence and activity between *D. melanogaster (mel)* and *D. simulans (sim)*, which diverged 1.4-3.4 million years ago^35^ (**Fig. 2c**), yet retain a similar ventral trichome pattern^12,36^. To this end, we aligned the *mel* and *sim* 266bp core *E3N* sequences, and identified five mutational changes (labeled A-E, **Fig. 2a**). Despite these differences, the overall *svb* expression pattern in the late-stage embryo is broadly conserved (**Fig. S2d-f**). However, the *sim E3N* activity is significantly weaker than that of the *mel E3N* in a *D. melanogaster* transgenic background, as the *sim E3N-lacZ* resulted in lower reporter expression and fewer nuclei with reporter expression (nuclear β-gal intensity, p = 1.43 x 10^-8^, number of β-gal positive nuclei, p = 3.03 x 10^-5^, **Fig. S2c**). This divergence can be explained by evolutionary compensation from other (“shadow”) *svb* enhancers, or evolutionary divergence of the trans-environment.

**Figure 2.**
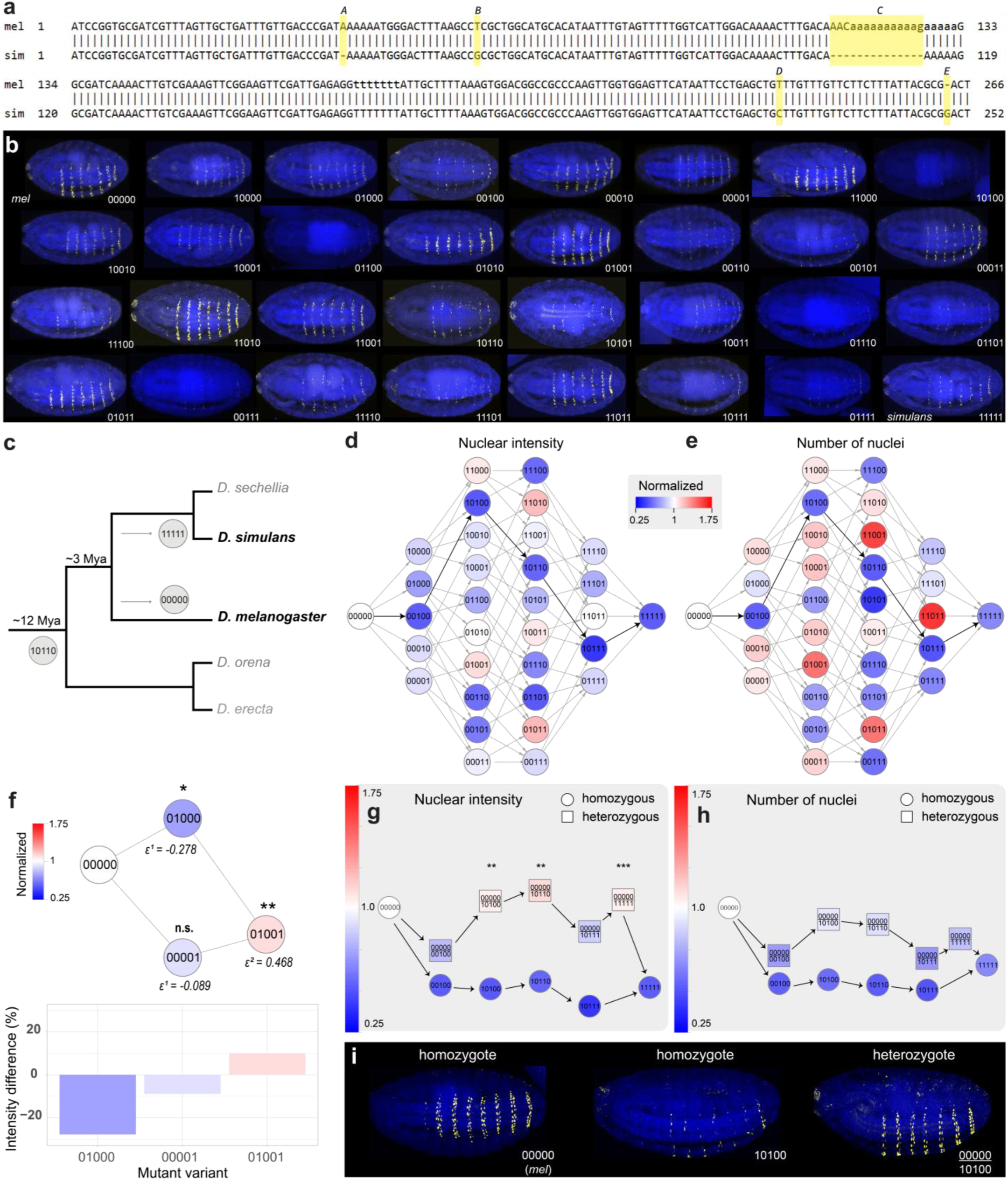
Higher-order epistasis and dominance in the evolutionary trajectory of the *E3N* enhancer. **(a)** An alignment of a minimized version of the modern-day *D. melanogaster E3N* (266 out of 292 bp) and modern-day *D. simulans* (ABCDE). Highlighted in yellow the five variations (titled A-E) in the genetic sequences. **(b)** Representative embryos for every line in the combinatorial *E3N* library showing the differences in the striped gene expression pattern of the lacZ reporter. **(c)** Evolutionary tree of the *melanogaster* species subgroup showing the changes in the *E3N* enhancer since the divergence of *D. melanogaster* and *D. simulans* 1.4-3.4 Mya, and the divergence of *erecta* complex 3.4-12 Mya. Here, ACD is the most likely ancestral version of the enhancer given the modern-day sequence of *D. erecta*. (**d-e**) Combinatorial evolutionary paths showing the possible intermediate forms from the genetic sequence of melanogaster (*mel*) to simulans (*sim*). Shown colors represent the average nuclear intensity (d) or the number of nuclei with enhancer activity **(e)** per ∼10 isogenic embryo normalized to the mean of the minimized melanogaster *E3N-lacZ* expression pattern. **(f)** Example of epistasis in the evolutionary trajectory of *E3N.* Variant *01000* has a significant negative effect on the nuclear β-gal intensity of *mel* (*00000*) (ε^1^ = -0.278, p = 0.016), whereas variant *00001* has no significant effect (ε^1^ = -0.089, p = 0.132) on *mel*. When the two variants are combined, there is a positive epistatic effect on the nuclear β-gal intensity of *mel* (ε^2^ = 0.468, p = 0.005). **(g)** A comparison of one of the evolutionary trajectories from modern-day *mel* to modern-day *sim* when every step is homozygous (represented as circles) versus heterozygous (squares), where one allele remains *mel* up until the last step to *sim*. The nodes in these paths are positioned on a y-axis relative to their normalized average intensity of the nuclei with enhancer activity. Linear models were fit independently for each cross that produced heterozygotes. Dominance deviations, defined as a significant p-value for βd, are marked with asterisks (* for p < 0.05, ** for p < 0.01, *** for p < 0.001). All non-significant heterozygote nodes are assumed to fit an additive model (βa). See Table S2 for a full description of the linear models. **(h)** The same evolutionary trajectory as in (g), but here the nodes in these paths are positioned on a y-axis relative to their normalized number of nuclei with enhancer activity. **(i)** Representative embryos for the homozygous modern-day melanogaster *00000*, the homozygous *10100* variant, and the heterozygous offspring of these two variants (*00000/10100*).

To disentangle these two hypotheses, we reconstructed the most probable sequence of the minimal *E3N* enhancer in the *melanogaster-simulans* ancestor by comparing the modern-day *E3N* sequences of *D. melanogaster* and *D. simulans* with the *E3N* sequence of *D. erecta* (3.4 - 12 Mya divergence from the *melanogaster-simulans* common ancestor^35,37,38^ (**Fig. 2c**, **Table S1**). We then hybridized *D. melanogaster* homozygous for this ancestral *E3N* enhancer with a *lacZ* reporter gene, with a reporterless *D. simulans* (**Fig. S2g-j**). We did not find differences in the reporter gene expression patterns for a single copy of the ancestral enhancer in a melanogaster embryo versus a hybrid embryo (**Fig. 2g-j**). This finding suggests that additional cis-regulatory changes in the *svb* locus, possibly in other enhancers that drive ventral *svb* expression, may have accompanied the changes in the *E3N* trajectory in either *D. simulans* and/or *D. melanogaster*.

### A complete genotype-phenotype map of *E3N* between *D. melanogaster* and *D. simulans* reveals extensive epistasis

By creating a genotype-phenotype map of the intermediate genotypes between the *mel* and *sim E3N* enhancer, one can trace all the possible evolutionary paths between the two sequences. To do this, we created a combinatorial mutagenesis library consisting of 32 enhancer variants representing every combination of the five genetic variations between the modern-day *mel (00000)* and *sim (11111) E3N* sequences (**Fig. 2a**), with each of the five sites either from *mel* (0) or *sim* (1). As an example, sequence *01000* contains the 266bp *mel* sequence, with a single T>G conversion at position 61 (**Fig. 2a,b**). These sequences were cloned into the *lacZ* reporter and integrated as described before (**Fig. 2b**). For each enhancer variant we quantified the nuclear β-gal intensity (**Fig. 2d**) and the number of nuclei with any *lacZ* expression (**Fig. 2e**) within the stripe pattern, and normalized all to the intensity of the *mel (00000)* construct. We inferred the effect of one, or multiple mutations in these variants by calculating first (ε^1^), second (ε^2^) or higher-order epistatic interactions (ε^n^)^39^ (see **Methods**).

We observe large differences in the number of nuclei with *E3N* activity between different constructs; among all single mutational events, *C (00100)* has the strongest effect on *mel (00000)* (ε^1^ = -0.476, p = 1.25 x 10^-6^ for the nuclear intensity, ε^1^ = -0.463, p = 5.12 x 10^-6^ for the number of nuclei). We also observe several instances of pairwise, and even higher-order epistatic interactions between the different combinations of mutations (**Fig. 2f**, see **Table S2**), which has been suggested to constrain enhancer evolution^40^. For example, construct *01000* has a significant negative (ε^1^ = -0.278, p = 0.016) or no effect (ε^1^ = -0.091, p = 0.456) on the nuclear intensity and the number of nuclei, respectively, and construct *00001* has no significant effect on both parameters (ε^1^ = -0.089, p = 0.132, and ε^1^ = 0.066, p = 0.373) (**Fig. 2f**). The combination of the two variants, construct *01001*, exhibits significant positive epistasis for both the nuclear intensity (ε^2^ = 0.468, p = 0.005) and the number of nuclei (ε^2^ = 0.435, p = 0.018) (**Fig. 2f**).

Across the evolutionary landscape of *E3N*, many mutational combinations markedly alter enhancer activity and display extensive epistasis, in stark contrast to the limited natural variation and conserved regulatory output observed in modern *D. melanogaster*.

### Heterozygosity expands possible evolutionary trajectories

We therefore asked how the evolutionary landscape for a fragile enhancer might be navigated via heterozygous intermediates. To investigate this, we mapped two hypothetical scenarios using the combinatorial *E3N* library (**Fig. 2g,h**). In the first scenario (“homozygous path”), mutations accumulate in a step-wise manner, but are homozygous at each step from *mel (00000)* to *sim (11111)*. In the second scenario (“heterozygous path”), mutations accumulate on one enhancer allele while the other allele remains *mel* until the final step to *sim*. Along the heterozygous path, we observed that intermediate heterozygotes often retained high enhancer activity despite carrying mutations on one allele that decrease expression in a homozygous context (**Fig. 2g**).

We observed dominance effects for three out of the five heterozygous combinations of alleles (See **Table S2**); here, the dominance of the *mel* allele resulted in higher enhancer activity than expected under an additive model (**Fig. 2g,h**). Moreover, except for the first step in this trajectory (from *mel*/*mel* to *mel*/*00100*), all subsequent heterozygous steps resulted in nuclear intensities statistically indistinguishable from those of the original homozygous *mel-lacZ*, but statistically different from its homozygous mutant counterpart. For example, the nuclear β-gal intensity of the heterozygous *mel/10100* offspring was statistically indistinguishable from the homozygous *mel* (p = 1), but differed significantly from homozygous *10100* (p = 6.99 x 10^-5^). In addition, we found a significant positive dominance deviation (βd = 0.288, p = 2.71 x 10^-3^), consistent with dominance of the *mel* allele in the heterozygote.

Using the combinatorial *E3N* library, we then reconstructed likely trajectories from the homozygous ancestral *10110* state to modern-day *mel* and *sim* sequences (**Fig. S2k**). In this scenario, one allele accumulates mutations while the other allele remains ancestral until the final steps towards the modern-day sequences of *melanogaster* and *simulans* (*00000* and *11111*) (**Fig. S2l**). Again, in an example of an evolutionary trajectory we found dominance effects in three of the five heterozygous steps, but here it was the mutation-accumulating allele exhibiting dominance over the stable allele.

To summarize, we observed dominance among heterozygous mutational steps that masks expression-reducing mutations in the evolutionary landscape of *E3N*.

### Enhancer dominance is cell-type specific

We then asked, could a wild-type allele also suppress pleiotropic expression caused by enhancer mutations? To this end, we focused on a (random) variant referred to as *145-2-lacZ* (**Fig. S3a)**, which exhibits ectopic enhancer activity in the wing and haltere primordia. *145-2-lacZ* also exhibits reduced *E3N* activity in its native stripe region nuclei (**Fig. 1m**, **Fig 3c**). We quantified β-gal intensity and the number of nuclei within the ventral stripe pattern for the homozygous wildtype *E3N* (**Fig. 3a**), homozygous *145-2* mutant (**Fig. 3b**) and their heterozygous offspring (**Fig. 3c**), as well as intensity in the the wing and haltere discs. For the wing and haltere primordia, we observed a strong additive effect (β_a_ = -1538.13, p < 2 x 10^-16^), and the intensity of the heterozygote discs differed significantly from both parental genotypes (p = 3.92 x 10^-05^ vs. homozygous E3N-lacZ and p = 4.00 x 10^-14^ vs. homozygous 145-2-lacZ, **Fig. 3d** and **Fig. S3f**). We found a significant negative dominance deviation (β_d_ = -781.0, p = 1.1 x 10^-6^), suggesting partial dominance of the *E3N-lacZ* allele in the cells that make up the wing and haltere discs.

**Figure 3.**
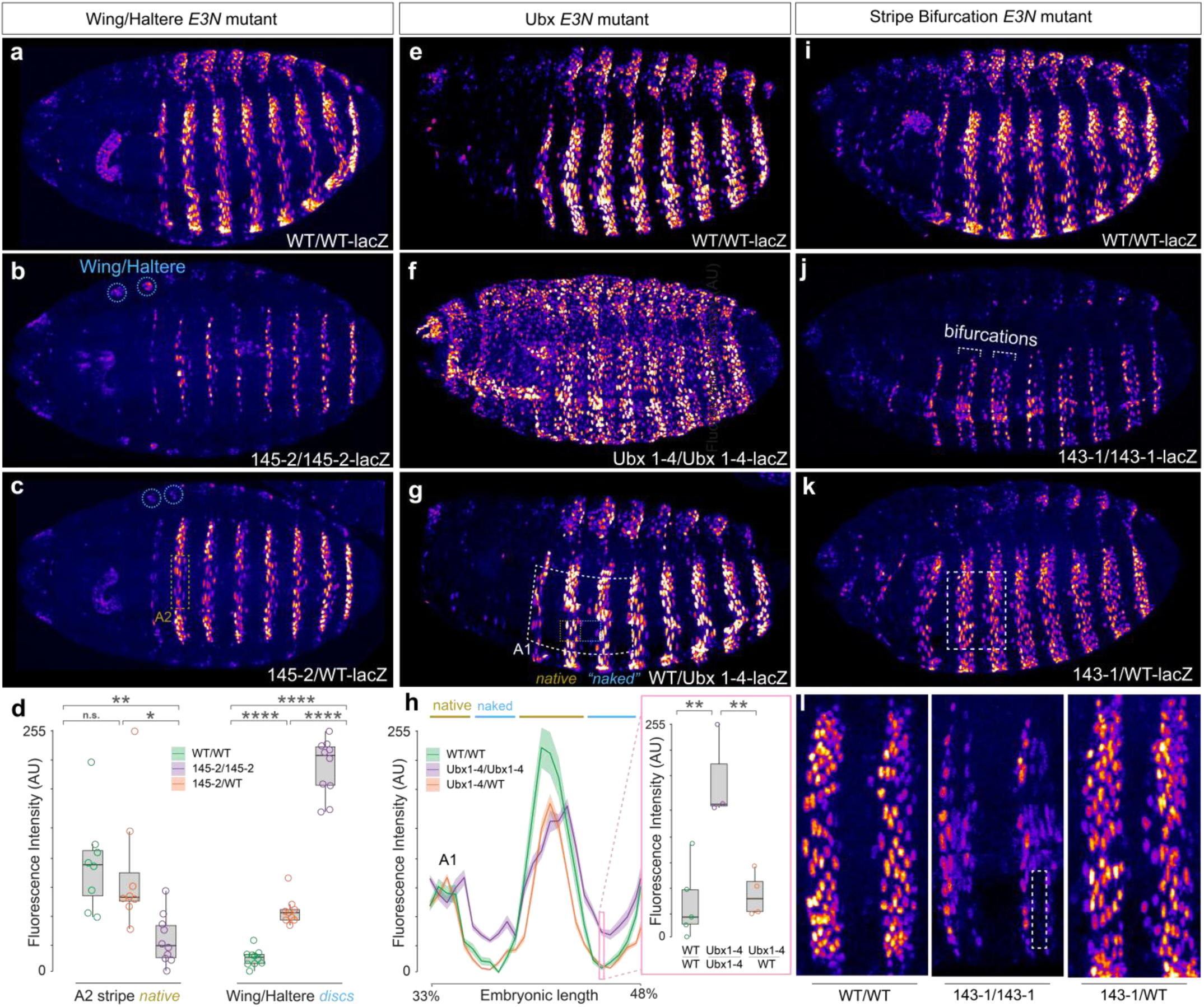
Cell type-specific regulatory dominance promotes robustness and innovation. **(a)** Representative image of a homozygous wildtype *E3N-lacZ* embryo at stage 15. Activity of the wildtype *E3N* enhancer does not result in *lacZ* expression in the wing and haltere disc. The embryo is stained with a β-gal antibody against the *lacZ* reporter protein β-gal. **(b)** Representative image of a homozygous mutant *E3N 145-2-lacZ* embryo at stage 15. Marked in blue circles are the wing and haltere discs where *lacZ* expression was measured across embryos. The embryo is stained with a β-gal antibody against the *lacZ* reporter protein β-gal. **(c)** Representative image of a heterozygous *E3N-lacZ/145-2-lacZ* embryo at stage 15. Marked in a mustard-yellow box is the region of abdominal stripe 2 (A2) where β-gal reporter intensity was measured across embryos. Marked in blue circles are the wing and haltere discs where β-gal reporter intensity was measured across embryos. The embryo is stained with a β-gal antibody against the *lacZ* reporter protein β-gal. **(d)** Boxplots of the mean fluorescence intensity of the β-gal reporter within the A2 stripe, and the wing and haltere discs, haltere disc of different embryos of the *145-2 E3N* variant at stage 15. Pairwise comparisons were performed using Tukey’s HSD test (confidence level 0.95). For the A2 stripe native boxplot, two out of three group comparisons were significant: WT/WT vs. 145-2/WT, p = 0,926; WT/WT vs. 145-2/145-2, p = 0,006; 145-2/WT vs. 145-2/145-2, p = 0.016. For the Wing/Haltere discs boxplot, all group comparisons were significant: WT/WT vs. 145-2/WT, p = 3,92 × 10^-5^; WT/WT vs. 145-2/145-2, p = 3 × 10⁻¹⁴; 145-2/WT vs. 145-2/145-2, p = 4 × 10⁻¹⁴. **(e)** Representative image of a homozygous wildtype *E3N-lacZ* embryo at stage 15. The embryo is stained with a β-gal antibody against the *lacZ* reporter protein β-gal. **(f)** Representative image of a homozygous *E3N Ubx1-4-lacZ* embryo at stage 15. The embryo is stained with a β-gal antibody against the *lacZ* reporter protein β-gal. **(g)** Representative image of a heterozygous *E3N Ubx1-4-lacZ/E3N-lacZ* embryo at stage 15. Region measured in (j) is depicted as a white rectangle from abdominal stripe 1 to 4 (A1-A4). Examples of a *native* (mustard yellow) and a *naked* (blue) region are shown within the ventral region. The embryo is stained with a β-gal antibody against the *lacZ* reporter protein β-gal. **(h)** Line graph showing the average of the β-gal reporter intensity profiles of multiple embryos for the genotypes depicted in (e-g) measured in the white box (stripe A1-A4). Shaded ribbons represent the standard error per genotype. Zones corresponding to *native* (mustard yellow) and *naked* (blue) regions in the wildtype pattern can be seen at the top of the figure. A boxplot of the fluorescence intensity at a position in the naked zone is shown in a pink box. Pairwise comparisons were performed using Tukey’s HSD test (confidence level 0.95), where two out of three group comparisons were significant: WT/WT vs. Ubx1-4/WT, p = 0,943; WT/WT vs. Ubx1-4/Ubx1-4, p = 2,79 x 10^-3^; Ubx1-4/WT vs. Ubx1-4/Ubx1-4, p = 5,62 x 10^-3^. **(i)** Representative image of a homozygous wildtype *E3N-lacZ* embryo at stage 15. Activity of the wildtype *E3N* enhancer results in uniform, lateral stripes of *lacZ* expression along the anterior-posterior axis of the ventral side of the embryo. The embryo is stained with a β-gal antibody against the *lacZ* reporter protein β-gal. **(j)** Representative image of a homozygous *E3N 143-1-lacZ* embryo at stage 15. In the *143-1 E3N* mutant, individual stripes exhibit bifurcation, where a single stripe splits into two parallel domains of *lacZ* expression (highlighted in white). The embryo is stained with a β-gal antibody against the *lacZ* reporter protein β-gal. **(k)** Representative image of a heterozygous *E3N-lacZ/145-2-lacZ* embryo at stage 15. Marked in a mustard-yellow box is the region of abdominal stripe 2 (A2) where β-gal reporter intensity was measured across embryos. Marked in blue circles are the wing and haltere discs where *lacZ* expression was measured across embryos. The embryo is stained with a β-gal antibody against the *lacZ* reporter protein β-gal. (**l**) Close-up of the stripe pattern of the different variants, where the lack of β-gal expression in the middle of the stripe (“bifurcation”) is highlighted with a white dotted box in the *143/143* variant.

By contrast, in the abdominal stripe 2 (A2), which is part of the native activity pattern of *E3N*, the heterozygote was statistically indistinguishable from the homozygous *E3N-lacZ* variant (p = 0.926), but significantly different from the homozygous *145-2-lacZ* mutant (p = 0.016) (**Fig. 3d**, **Fig. S3f,g**). Thus, the heterozygous *E3N* activity in the cells of the ventral stripe resembles the wildtype *E3N-lacZ* activity pattern, consistent with dominance of the wildtype allele, although the dominance-deviation test itself was not significant (β_d_, d = 664.9, p = 0.157).

To confirm that this apparent interallelic repression was not unique to the ectopic Wing-Haltere phenotype, we also crossed the wildtype *E3N-lacZ* enhancer with an *E3N* variant called *Ubx1-4-lacZ* (**Fig. S3b)**, which exhibits a severe reduction of reporter expression at stage 15^41^, and an increase in ectopic expression between the stripes at stage 16 (**Fig. 3f**). We created heterozygous offspring of *Ubx1-4-lacZ* with the wildtype *E3N-lacZ* (**Fig. 3e)**, and quantified the β-gal reporter intensity of the stripe pattern at stage 16. Surprisingly, we did not observe any ectopic *lacZ* expression in the “naked” regions of the embryo in the heterozygote, suggesting that within these cells the mutant ectopic expression is fully repressed by the presence of the wildtype *E3N* allele in a dominant manner (**Fig. 3h**).

Lastly, we crossed the wildtype *E3N-lacZ* line with another *E3N* variant called *143-1* (**Fig. S3c)**, which creates a “bifurcated” stripe pattern where a normally single band splits into two parallel rows of nuclei with expression (**Fig. 3j**). We found that the heterozygous expression pattern is similar to the wildtypes embryos’, where a single peak is observed within stripe A3 (**Fig. 3i-l** and **Fig. S3d**). This again suggests that the loss of *E3N* activity in the middle of a single band in the mutant embryo is rescued by the presence of a wildtype *E3N* allele in a dominant manner.

In summary, through multiple examples we demonstrated that a wild-type enhancer allele in heterozygous conditions can exert a dominant regulatory influence: it consistently represses aberrant ectopic expression domains and rescues deficits in normal expression caused by the mutant allele.

### Enhancerless and promoterless E3N alleles complement in trans

Classically, transvection is defined as a proximity-dependent interaction between homologous alleles. We therefore tested whether the regulatory dominance we observed similarly depends on genomic distance. We inserted the *E3N-lacZ* construct at the *attP40* landing site on chromosome 2, which recapitulates the expression pattern observed at the chromosome 3 insertion site (**Fig. S3h-j**). When the wild-type allele on chromosome 2 was combined with the mutant *145-2* allele on chromosome 3, enhancer activity was significantly reduced relative to heterozygotes in which both alleles were inserted at the same locus (**Fig. S3j**). Thus, the buffering effect of the wildtype allele is impaired when located on a different chromosome. Notably, this configuration resulted in underdominance: the heterozygote with alleles on different chromosomes exhibited significantly lower enhancer activity than either homozygous wildtype or mutant genotypes (β_d_ = -1176.0, p = 1.53 x 10^-5^).

We next tested whether the homologous enhancers can act on a promoter in trans. We compared enhancer activity from a single *E3N-lacZ* allele (“E3N-lacZxattP2”), to that of the same allele in a heterozygous context paired with an identical construct in which the promoter was replaced by a neutral spacer (“E3N-lacZxspacer”, **Fig. S4a,b**). The spacer construct alone does not drive reporter expression (“spacer”, **Fig. S4a,b**). Although hemizygous *E3N-lacZ* (“E3N-lacZxattP2”) tended to have a lower activity than when paired with the spacer allele, the confidence intervals overlap, indicating that the difference is not statistically robust.

We then asked whether functional complementation occurs when enhancer and promoter are separated across homologs. An allele lacking the enhancer (ΔE3N) was combined with a homologous allele lacking the promoter (“spacer” or “Δpromoter”), neither of which drives expression independently (**Fig. S4c,d**). Notably, we observed that more expression is restored without the insertion of a neutral spacer sequence at the deleted promoter site (“Δpromoter”).

Because promoters differ in their ability to transvect and in their preference for regulation in cis or trans^24,42^, we replaced the *hsp70* promoter with the native *svb* promoter (“E3N-svbp-lacZ”) and repeated the spacer assay. Pairing *E3N-svbp-lacZ* with the promoterless spacer allele again increased enhancer activity relative to the hemizygous condition (**Fig. S4e,f**), indicating that interallelic interactions are not specific to the *hsp70* promoter. Interestingly, the hemizygous *E3N-svbp-lacZ* exhibited fewer β-gal–positive nuclei than expected from simple additivity relative to its homozygous parent, a deviation not observed with the hsp70 promoter construct.

Together, these results demonstrate functional interallelic complementation consistent with a proximity-dependent, transvection-like mechanism underlying the dominance effects observed at *E3N*.

### Visualing the transvection hub

To understand the mechanism behind the different degrees of dominance in different cell types, we employed a two-color reporter assay with fluorescent in situ hybridization. Heterozygous embryos were generated carrying the *WT E3N* enhancer variant driving a *dsRed* reporter gene, and the *145-2 E3N* variant driving a *lacZ* reporter (**Fig. 4e**). We then performed in situ Hybridization Chain Reaction (HCR) for the two reporter genes in the heterozygous *145-2-lacZ/WT-dsRed*, and simultaneously stained for the transcription factor Ubx (**Fig. 4a,c-d**), as active transcription sites of *svb* are known to colocalize with local high concentrations of Ubx^6^.

**Figure 4.**
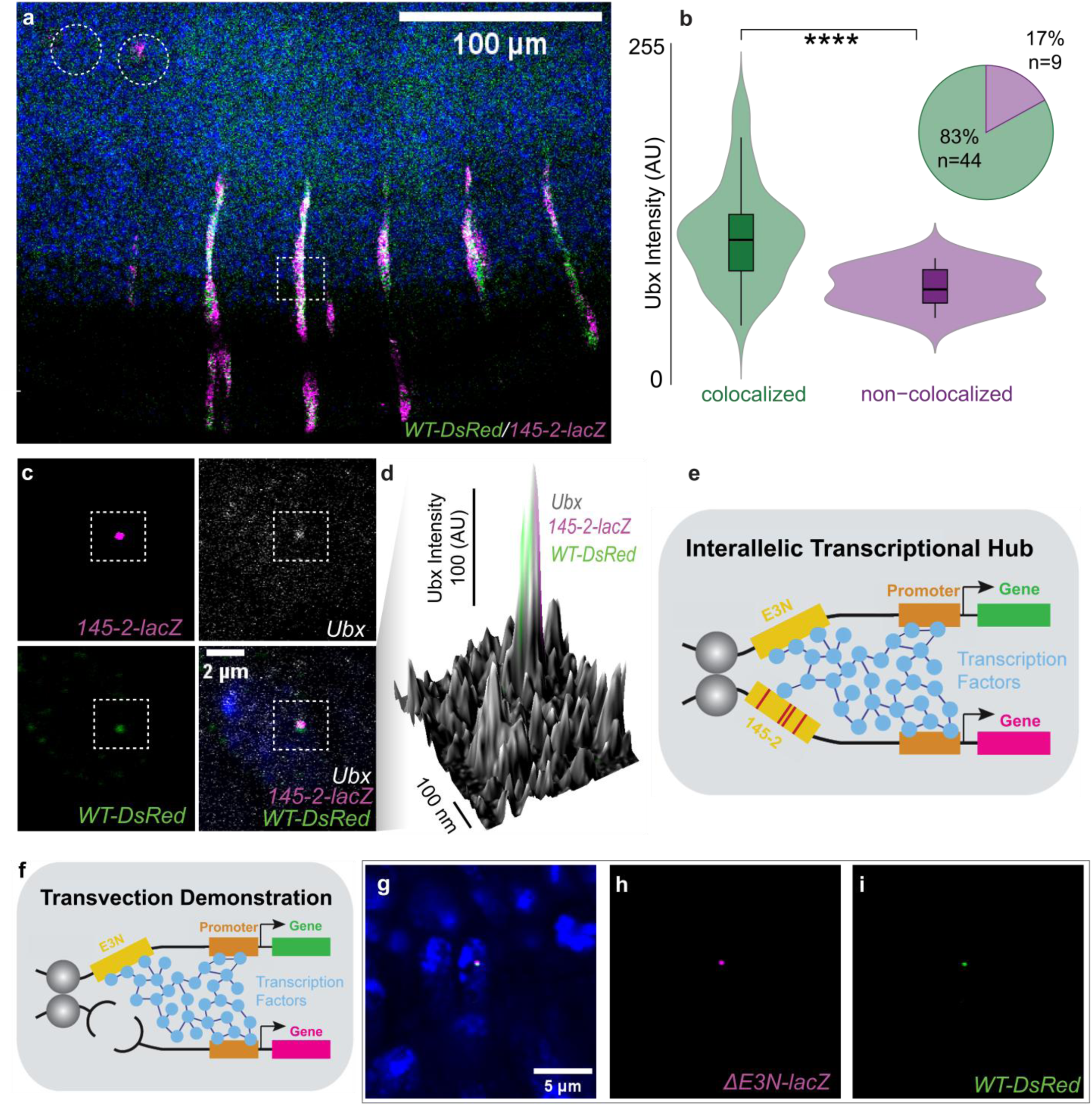
Interactions between homologous trans-alleles coincide with transcriptional transvection hubs. **(a)** Stage 15 embryo resulting from a cross of the wildtype *E3N-lacZ* line, integrated at *attP2* (chr.3L) with ectopic *E3N* mutant *145-2* driving *lacZ*, integrated at *attP2* (chr.3L). RNA of both the *lacZ* and *dsRed* reporter genes are labeled by HCR RNA-FISH. **(b)** Violin plot showing the difference in Ubx concentration between colocalized (green) and non-colocalized (purple) transcription sites of *lacZ* and *dsRed*. Colocalized sites exhibit significantly higher Ubx intensity compared to non-colocalized loci (p=1.2e^-5^; two-sided Student’s t-test). The pie chart summarizes the proportion of observations in each category. **(c)** Close-up of a nucleus (in blue, DAPI) in the ventral zone, highlighted by a dotted-striped rectangle in (a). Separation of the different channels reveals colocalization of *lacZ* and *dsRed* nascent RNA at a shared transcription site, accompanied by a local enrichment of Ubx protein. **(d)** 3D surface plot of Ubx protein levels from antibody staining of the nucleus pictured in (c). Ubx signal is shown in arbitrary units (AU) on a 0-255 scale, and presence of *lacZ* and *dsRed* signal is depicted in magenta and green, respectively. **(e)** Schematic of the interallelic transcriptional hub for the genotype described in (a). **(f)** Schematic of the construct used in g, where the two different reporter genes enable one to differentiate between the signals coming from the two alleles. Relative to panel (e), the *145-2* variant is removed. A homozygous construct with two alleles without enhancers shows minimal expression in the *Drosophila* embryo. (**g-i**) Overview of some nuclei (in blue, DAPI) within the ventral zone of a stage 15 embryo resulting from a cross of the wildtype *E3N-dsRed* line, integrated at *attP2* (chr.3L) with a construct without an enhancer, integrated at *attP2* (chr.3L). RNA of both the *lacZ* and *dsRed* reporter genes are labeled by HCR RNA-FISH.

Within homozygous *145-2-lacZ* embryos, we identified several active *lacZ* transcription sites at the time of embryo fixation in the wing and haltere discs (**Fig. S5f-i**). In contrast, in the heterozygous *145-2-lacZ/WT-dsRed* embryos, no active *lacZ* transcription sites in the discs were identified (**Fig. S5b-e**). As expected, no *dsRed* signal was detected in the wing and haltere discs of homozygous *WT-dsRed* embryos. Conversely, in heterozygous *145-2-lacZ/WT-dsRed* embryos we detected (mature) RNA molecules from both reporter alleles in both wing and haltere discs (**Fig. S5e**). This indicates that trans-homologous activation still occurs, albeit likely at a reduced frequency. Such reduced interallelic activation could explain the negative dominant effect exerted by the *WT-dsRed* allele on gene expression in the heterozygote within the wing and haltere discs (**Fig. 3d**).

In the nuclei in the ventral zone of the heterozygous embryos that contain active *lacZ* transcription sites, we observed colocalization with the *dsRed* transcription site in 83% of the cases (**Fig. S5a** and **Fig. 4b-d**). In addition, we observed a significantly higher concentration of Ubx protein around the *lacZ* transcription site when it was colocalized with *dsRed* (p = 1.2 x 10^-5^, **Fig. 4b**).

Lastly, we performed the two-color reporter assay on the heterozygous offspring of *ΔE3N-lacZ* and *WT-dsRed* embryos (**Fig. 4f**). As described before, homozygous *ΔE3N-lacZ* embryos exhibit minimal *E3N* enhancer activity (**Fig. S4c,d**). Here, we again observed colocalization of the *lacZ* and *dsRed* transcriptional sites (**Fig. 4g-i**).

All together, these results are consistent with a transvection-like interallelic hub contributing to the dominant effects we observe, where cell types that exhibit more interallelic interactions also display stronger dominance effects (**Fig. S6**).

## Discussion

Using a complete genotype–phenotype map between the *D. melanogaster* and *D. simulans E3N* enhancers, we show that most homozygous evolutionary paths exhibit reduced enhancer activity and (higher-order) epistasis. Such epistatic constraints are predicted to bias evolutionary trajectories and the order in which mutations can accumulate^43–45^. However, when the same mutational steps are in a heterozygous context, their phenotypic consequences are frequently masked by regulatory dominance. This buffering may relax constraints on enhancer evolution^12,34,46–48^, enabling trajectories that preserve transcriptional output despite individually deleterious intermediate states. Our results, therefore, highlight heterozygosity as a possible feature of diploid genomes that can mitigate constraints imposed by deleterious mutations and regulatory epistasis.

The dominance we observe is mechanistically distinct from classical explanations based on protein dosage^49^, network feedback^50^, or trans-acting variation^51^. Instead, dominance arises solely from differences in enhancer sequence and depends on the physical proximity of homologous alleles. We show that this buffering is impaired when alleles are placed on different chromosomes and that enhancerless and promoterless alleles can complement one another in trans. Together with allele-specific transcription imaging, these results support a model in which homologous enhancers and promoters participate in shared transcriptional hubs^23,30^, allowing regulatory information to be pooled across alleles. In this framework, dominance emerges as a property of regulatory architecture rather than of gene products, extending the classical concept of transvection^52^ from a genetic phenomenon to a more general mode of regulatory integration.

Crucially, interallelic dominance is not uniform across tissues. In heterozygotes carrying pleiotropic enhancer mutations, wild-type alleles can fully repress ectopic activity in some cell types while permitting mutant-driven expression in others. Multiple factors may underlie these observations, such as cell-type specific differences in interallelic pairing strengths^53^, chromatin accessability, post-transcriptional regulation^54^ and competition for TFs^55,56^. We also speculate that the exact composition of TFs in an interallelic hub may influence the degree of dominance, where certain cell-type-specific TFs may promote transcriptional condensation^29,57^, and others may quench the activity of local activators^58^. This cell-type-specific dominance creates a mosaic regulatory outcome in which robustness and novelty coexist within the same organism. Such modular buffering resolves the tension in regulatory evolution: how mutations in essential, pleiotropic enhancers can persist long enough to contribute to innovation. By compartmentalizing their effects, interallelic hubs may allow mutations to be selectively exposed in permissive contexts while remaining buffered in developmentally critical tissues.

Our findings support a general model in which interallelic gene regulatory hubs can act as evolutionary stepping stones by distributing dominance relationships across cell types and regulatory contexts. Rather than having a single, uniform phenotypic effect, each enhancer allele can experience a distinct dominance relationship depending on cellular environment and developmental stage (outlined in **Fig. S6**). This added dimensionality could expand the range of accessible evolutionary trajectories. Extensive somatic homolog pairing has predominantly been characterized in Diptera and budding yeast^59,60^, but pairing-associated interallelic interactions have more recently been reported in maize and pigs as well^61,62^. Moreover, interallelic and interchromosomal transcriptional hubs are increasingly recognized across metazoans^21,23,25–27,63^, suggesting that analogous regulatory principles may operate more broadly. Indeed, analyses of heterozygous regulatory variation in humans, including dominance eQTL and allele-specific expression studies, reveal frequent departures from additivity and strong tissue dependence of cis-regulatory effects^64,65^.

Together, these findings identify cell-type-specific interallelic transcriptional hubs as a possible feature of diploid genomes that reconciles robustness and evolvability; gene expression is maintained in the face of potentially disruptive mutations (robustness), while concurrently allowing the exploration of new phenotypic effects in a cryptic or compartmentalized way (innovation). By linking dominance to regulatory architecture^66^ and three-dimensional genome organization^67,68^, this work provides a framework for understanding how seemingly fragile cis-regulatory elements preserve their function yet remain evolutionarily flexible.

## Material and Methods

### Reagents

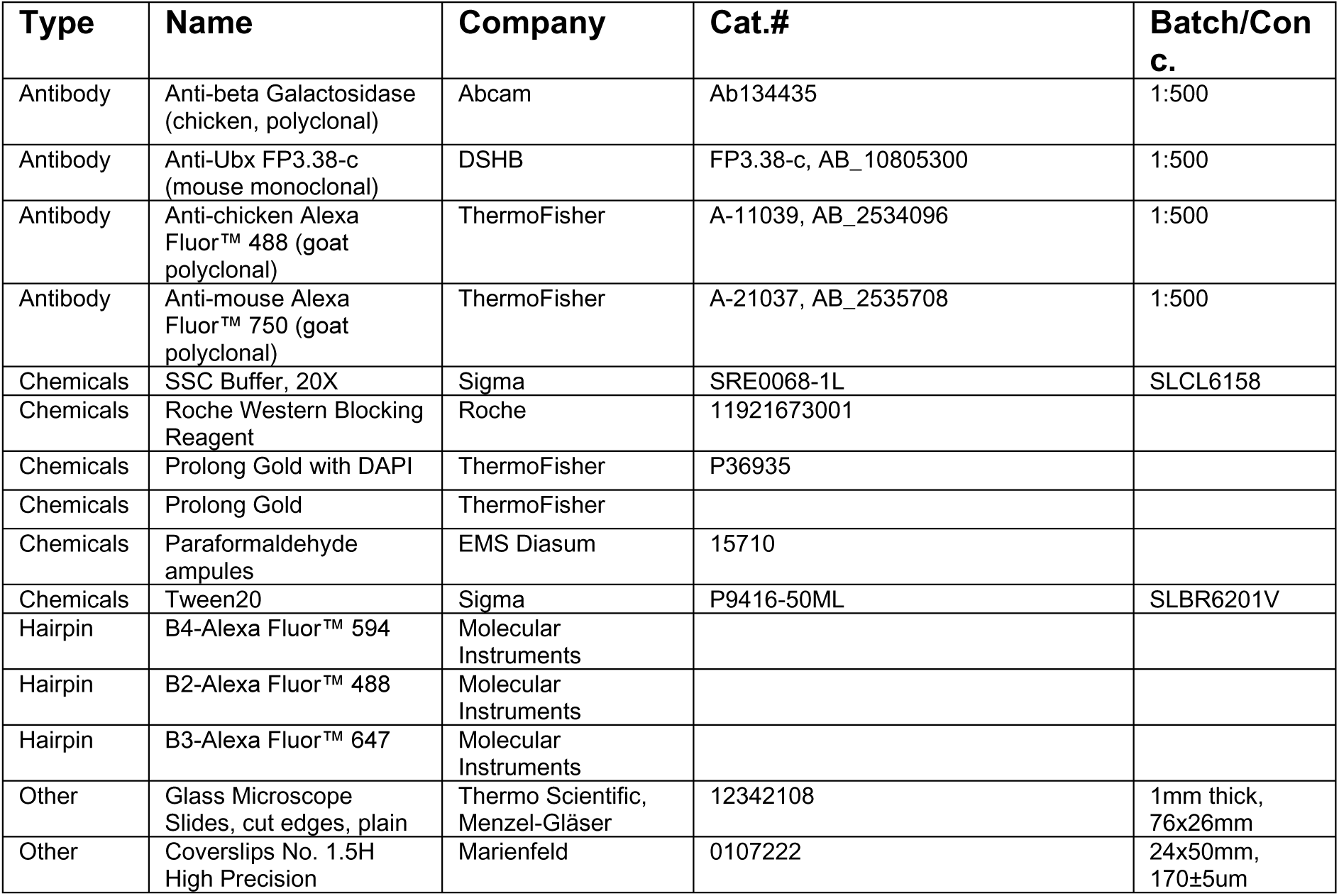

### Fly husbandry

Crosses were performed under standard conditions at 22℃ or 25℃ in plastic vials supplemented with fly food (1.8% m/v yeast, 8.5% m/v cornmeal, 0.7% m/v agar, 6.3% m/v sugar beet syrup, 0.5% v/v propionic acid, 0.9% v/v nipagin 20% in ethanol). Transgenic lines were created using the *attP2* (chromosome 3L) or *attP40* (chromosome 2) lines (BDSC# 25710 and 36304, respectively). Outcrossing was performed using the w1118 line (BDSC# 5905). *D. simulans* (gift from Nicolas Gompel) and *D. melanogaster Ind* (“Mysore” strain, old stock #3114.4 from National Drosophila Species Stock Center) lines were used to compare *svb* gene expression patterns. FlyBase was used to find information on different stocks^69^.

### *E3N* SNP identification

To identify the SNPs in the *E3N* sequence, *D. melanogaster* sequences from the Nexus V1.1 database were used^33^. First, duplicate identifiers, and sequences containing >10% of masked nucleotides (N) were removed, which are sites within 3bp of a called indel, or large genomic intervals of apparent heterozygosity. The X-chromosome of the remaining 821 sequences was aligned in JalView^70^ (v2.11.1.3) using the MUSCLE (multiple sequence comparison by log-expectation, v3.8.31) algorithm^71^, and the *E3N* sequence was cut out of this alignment and realigned (see **File S1** for the alignment). Next, SNPs were identified using SNP-sites^72^ (v2.5.1), and the occurrence of certain nucleotides was established using the R package SeqinR^73^ (v4.3-4). SNPs were called if they occurred at a frequency of ≥1%.

### *E3N* combinatorial library generation

We created an *E3N* sequence alignment between *D. melanogaster,* the sister species *D. simulans,* and the outgroup species, *D. erecta* using MUSCLE^71^, MAFFT^74^, and ClustalO^75^ (**Table S1**). These alignments revealed a 266 bp, semi-conserved, core region between the three paralogs, which contains 5 differences between the *melanogaster* and *simulans* homologs. We created a “clipped” version of *E3N* that contains the semi-conserved, 266 bp sequence. In *melanogaster*, this clipped sequence lacks the first 11 bp and the last 15 bp from the original *E3N* construct^31^. We then created a combinatorially complete library (N=2^5^=32 constructs, including the modern-day sequences of *melanogaster* and *simulans*) that contains all possible mutational steps that could occur between the semi-conserved *melanogaster* and *simulans* homologs (**Table S1**).

### Fly transgenesis

Enhancer sequence synthesis and fly transgenesis was performed as described previously^34^. Variants of the *E3N* enhancer were designed and cloned into the HindIII (5’) and XbaI (3’) sites of the placZattB reporter plasmid (GenScript), and integrated into the *attP2* and/or *attP40* integration sites in D. melanogaster genome using the PhiC31 integrase system as performed before^12,76^. As described^34^, the subset of 91 mutant lines from the original 749 *E3N* mutant variants was selected using a random module; 10 lines containing one mutation, 9 with two mutations, 9 with three mutations, 9 with four mutations, 7 with five mutations, 9 with six mutations, 9 with seven mutations, 10 with eight mutations, 8 with nine mutations, and 10 with ten mutations. Similarly to the placZattB reporter plasmid, variants of the *E3N* enhancer were cloned into a dsRedattB reporter plasmid^21^,and integrated at both *attP2* and *attP40*. The svbp, spacer sequence and Δpromoter variants^77^ of the placZattB constructs were integrated at *attP2*. See **Table S1** for all the sequences used in this study.

### Embryo collections and fixations

Flies were put in egg-laying cages four days before collection to acclimate, and fixations were performed after overnight collections (approximately 15 hours) at 25℃. Embryos were manually fixed as described before^12^, and kept at -20℃ in methanol until further processing.

### Reporter gene expression assays

#### HCR in situ hybridization

Hybridization Chain Reaction (HCR) in situ hybridization was performed using probes and hairpins synthesized by Molecular Instruments (MI). The protocol for whole-mount Drosophila embryos^78^ was used, adapted for split-initiator probes (v3.0)^79^. As described before^80^, several changes were made to the protocol: no treatment with ethanol, xylene, and proteinase K at the start, a reduction of the percentage of dextran sulphate in the probe hybridisation and amplification buffers to lower the viscosity, and the addition of a postfixation step at the end of the protocol to stabilise the signal. For the *shavenbaby* expression pattern the embryos were incubated with the *svb* probe overnight (16 hours), which, after washing, was followed by an overnight amplification step (16 hours).

#### Immunofluorescence staining

Immunofluorescence was performed as described before^34^, with an overnight primary chicken anti-beta Galactosidase antibody staining (1:500, abcam) at 4℃ and a two-hour secondary goat anti-chicken Alexa Fluor™ 488 antibody staining (1:500, ThermoFisher) at room temperature. Embryos were mounted on slides with Prolong Gold with DAPI (ThermoFisher) and left to cure for three days in the dark at room temperature before imaging.

#### HCR in situ hybridization with immunofluorescence staining

To detect active transcription sites of the *lacZ* (Alexa Fluor™ 594) and *dsRed* (Alexa Fluor™ 488) reporter genes, the amplification time in the HCR protocol described above was brought down to 60 minutes at room temperature. HCR was combined with immunofluorescence^80^, where after the postfix step at the end of the HCR protocol, the embryos were washed with SSCT (5xSSC, 0.1% Tween20), incubated for 25 minutes with SSCT blocking solution (Roche Western Blocking Reagent #11921673001; diluted 1:5 in SSCT), and then incubated overnight with mouse anti-Ubx (1:20 in SSCT blocking solution, FP3.38-c, DSHB, supernatant) at 4℃.

### Image acquisition

For the immunofluorescence-only conditions, images were acquired on a Zeiss LSM880 confocal microscope using the semi-automated adaptive feedback 3D microscopy pipeline^81^. In short, low-resolution overview tile scans were used to manually identify embryos of the correct orientation and developmental stage. For each selected embryo, the center and orientation is identified in 3D through a rapid scan, which is used to guide the acquisition of a high-resolution multichannel stack through the entire embryo. Final images were acquired in 16-bit depth at 1024x1024 resolution, with a a 50.4um pinhole, Z-step size of 1.404um, and a total Z-depth of 42.127um (30 Z steps), using a 20x/0.8 NA air objective. The complete imaging pipeline, including the repository, setup details, and a quick-start guide, is available at: https://git.embl.de/grp-almf/feedback-fly-embryo-crocker

For the samples processed with HCR RNA-FISH combined with immunofluorescence, images were acquired on a Leica Stellaris confocal microscope equipped with a White Light Laser (WLL) and HyD detectors. Each embryo was first imaged at low magnification (20x/0.75 NA oil objective, HC PL APO CS2) to obtain an overview, followed by high-magnification imaging (63x/1.40 NA oil objective, HC PL APO CS2) of different regions of the same embryo. High-resolution images were acquired in 16-bit depth at 2728x2728 resolution, with a 131.715 pinhole, bidirectional scanning (speed = 400), and a Z-step size of 0.412um. Total Z-depth varied between 15-17 steps depending on the region imaged. Sequential line scanning was used to acquire the following channels: Channel 1 (DAPI) – Diode 405 nm (UV light); Channel 2 (Ubx-AF750) – WLL 730 nm; Channel 3 (*dsRed*-488) – WLL 488 nm; and Channel 4 (*lacZ*-594) – WLL 590 nm. For every embryo, a single 63x image of the ventral region was used for subsequent quantitative analyses of individual nuclei.

### Image analysis

#### Image preparation

Images were preprocessed using the image analysis program Fiji^82^: images were max projected and cropped to fit only the full embryo. For the automated analysis with Cellpose^83^, images were manually cropped to only encompass the seven most posterior ventral stripes.

#### Automated image analysis pipeline

To study the effect of the mutations on the enhancer, we analyzed two components of the stripe pattern (average nuclear intensity and number of nuclei with intensity) using the pretrained Cyto3 model of Cellpose^83^, for which an automated image analysis pipeline can be found here: https://git.embl.de/grp-cba/fly-embryo-expression-pattern

#### Manual image analysis – different cell types

To extract the β-gal reporter intensity of different cell types within the 145-2 mutant version of the *E3N* line, manual image analysis was performed. For the wing and haltere disc, a circle with a radius of 15 pixels (7.784um) was centered around both of the discs, and raw intensity values were extracted in Fiji for multiple embryos for both discs. Then, intensity values were normalized to an arbitrary unit (AU) range of 0-255 and boxplots were generated in R (v4.2.2) using ggplot (v3.5.1) for the β-gal reporter intensity in the wing disc, the haltere disc, and the average of the wing and haltere discs.

For the intensity measurements of the A2 stripe zone, a rectangle was created with a height equal to the ventral nerve cord (VNC) of that embryo, and a width equal to the width of the *lacZ* stripe. Raw intensity values were extracted in Fiji for multiple embryos. Then, intensity values were normalized to an arbitrary unit (AU) range of 0-255 and boxplots were generated in R (v4.2.2) using ggplot (v3.5.1).

#### Stripe pattern profiles

Per genotype, montages of max projected embryos were created. A segmented line (width = 100 pixels) was drawn along the midline of the VNC in Fiji, starting at the anterior side of stripe A1, and ending at the posterior end of the embryo. Lines were added to ROI Manager, and raw intensity profiles per embryo were generated using the *Plot Profile* function. The profiles of embryos and genotypes contain pixel gray values along our region of interest and were combined into one CSV file for downstream analysis. The average profile of every genotype was generated in R (v4.2.2) using the dplyr (v1.1.2) and tidyr (v1.3.0) packages.

First, the positional coordinates (distance in microns) were normalized to a continuous scale from 0 to 1 for every embryo. To compare different embryos of the Ubx1-4 dataset, the normalized positions were binned at intervals of 0.01, and each bin was centered to represent its midpoint along the axis. Then, the mean intensity, standard deviation, and standard error of the mean were calculated for each bin in a genotype. To compare different embryos of the 143-1 dataset, a common x-axis grid of 1000 equally spaced points was defined, on which the intensity values were interpolated using linear interpolation. Intensity traces were slightly shifted along the normalized axis to align peak position with the reference trace. For visualization, the profiles were cropped to a encompass stripe A1 to A3 (Ubx1-4), or only stripe A3 (143-1), and mean intensity values were plotted as line graphs with shaded ribbons representing the standard error using the ggplot2 (v3.5.1) package. To compare fluorescence intensity across genotypes in the naked zone in the Ubx1-4 dataset, we first identified the bin in the naked zone between stripe A2 and A3 in which the wildtype average intensity reached its lowest value. For each embryo, all pixel values mapping to this bin were averaged, yielding a single intensity value per embryo, which were visualized in a boxplot.

#### Colocalization spot and hub analysis HCR

Because the amplification time of the HCR protocol was substantially reduced, signal amplification was limited. Under these conditions, large, bright nuclear foci were interpreted as sites of nascent transcription, whereas smaller, discrete puncta were interpreted as (near-)individual, or low-copy mature RNA molecules. Colocalization was established if two transcriptional spots were within 360nm of another^30^. In the mutant allele channel (*lacZ*), nuclei were identified containing a single transcriptional spot that was present in at least two Z slices (0.4um). Within each selected nucleus, the coordinates of the pixel with the maximum intensity per channel was identified using the *Find Maxima…* function, and the Euclidean distance between the wildtype and mutant transcriptional spots was calculated. Then, a circle with a radius of 170nm was centered on the pixel with the maximum intensity in the mutant channel and the average intensity per pixel, the minimum, and the maximum intensity were recorded in the mutant *lacZ* RNA, wildtype *dsRed* RNA and Ubx protein channel. In addition, for every 170nm measurement, the same measurements were performed in the entire nucleus of that spot for every channel, and for a randomly generated 170nm radius spot in that nucleus. This analysis was performed for approximately eight nuclei per embryo, and for seven embryos of the same developmental stage 15, which is the moment the gut gets a heart-shaped form.

To allow for comparison across embryos, intensity values were normalized to an arbitrary unit (AU) range of 0-255, where the minimum value for each channel recorded within an embryo became zero, and the maximum value became 255. Transcriptional spots were classified as *colocalized* if the distance between the pixel of maximum *lacZ* intensity and the pixel of maximum *dsRed* intensity was ≤360 nm, or if the average intensity per pixel for *dsRed* within the 170nm radius circle centered on the pixel with the highest lacZ intensity was at least twice the average intensity per pixel for *dsRed* across the entire nucleus. Spots within a nucleus not meeting either criterion were classified as *non-colocalized*. Statistical comparisons of the average Ubx intensity per pixel within a 170nm radius centered on the pixel of maximum *lacZ* intensity within the nucleus were performed between transcriptional spots classified as colocalized or non-colocalized using two-sided Student’s t-tests.

### Non-additivity analyses and statistics

#### Epistasis

The effect of one, or multiple mutations in the intermediate genetic variants in the combinatorial *E3N* library were inferred by calculating first (ε^1^), second (ε^2^) or higher-order epistatic interactions (ε^n^)^39^. Here, ω is defined as the nuclear intensity or the number of nuclei with reporter gene expression for every isogenic embryo inferred by an automated analysis pipeline using Cellpose^83^, relative to the *D. melanogaster* wildtype (00000):

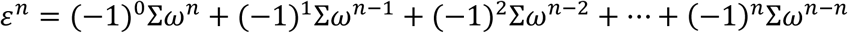

The propagated errors (*δσ*) associated with every genotype (*ω*[*g*]) were then calculated as follows:

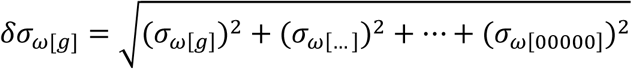

Here, “[…]” and “…” represent the different combinations of mutations that make up [g]. Next we performed a one-sample t-test using every epistatic term and its respective propagated error.

#### Dominance

To test whether the heterozygote within a genotype triplet (AA, AB, BB) significantly differed from its parents’ phenotype, we fitted a linear model for the triplet. Pairwise posthoc comparisons between genotypes were performed using the emmeans package (v1.11.2) with Tukey’s honestly significant difference (HSD) adjustment for multiple testing.

To test the deviation in the intensity and the number of nuclei within the reporter gene stripe pattern, we fit a linear model for each genotype triplet (AA, AB, BB) associated with every cross we did. Genotypes were rewritten into additive (a) and dominance (d) components:

- a = +1 for AA, 0 for AB, and -1 for BB
- d = 1 for AB, 0 for AA and BB

For both the intensity and the number of nuclei values that we generated using the Cellpose pipeline as described before, we modeled each cross as:

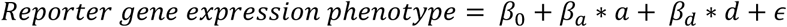

where *β_a_* represents the additive effect, and *β_d_* the dominance deviation. Overviews of the linear models, generated in R using the lm() function, can be found in **Table S2**.

#### Additional statistics

All statistical analyses were performed in R (v4.2.2). To determine whether each of the SNPs had a significant effect on the activity of the wildtype *E3N* enhancer, we fitted a one-way analysis of variance (ANOVA) with genotype as a fixed effect for both the reporter intensity and the number of nuclei with reporter intensity. Post hoc comparisons were performed using Dunnett’s multiple-comparison test. Adjusted p-values were obtained using the multcomp package (v1.4-28). For the variant that proved to be significantly different from the wildtype sequence, we calculated effect size using Hedges’ g with the effectsize package (v1.0.1). Estimates, standard errors, t-values, adjusted p-values and effect sizes can be found in **Table S2**.

For the evolutionary trajectories between mel/00000 and sim/11111, within-triplet differences were tested using linear contrasts in the emmeans package. P-values were adjusted using the Holm method.

#### Visualization

To generate the mutational landscapes, the igraph package (v2.1.4) in RStudio was used. Here, nodes represent binary mutation states and edges connect nodes that differ by a single mutation (a Hamming distance of 1). The mean intensity and number of nuclei with enhancer activity per genotype were normalized relative to the respective mel/00000 phenotypes. The normalized averages were mapped to a color gradient (from blue (0.3) to white (0.0) to red (1.25)) to visually encode relative expression levels across the network.

## Additional Supplementary Data

**Supplementary Figures.** S1-S6.

**Supplementary File 1**. Fasta alignment of the 821 sequences of E3N from the Nexus V1.1 database used for the SNP calling

**Supplementary Table 1**. Sequences and fly lines used in this study

**Supplementary Table 2**. Data used in this study

## Acknowledgements

We thank Matthew Benton and Alessandra Reversi from EMBL’s *Drosophila* transgenesis service for fly embryo injections to generate the transgenic lines. We also thank Lautaro Gandara, Paco Majic and Santiago Herrera-Álvarez for their advice and support during this project, and Albert Tsai for helpful discussions regarding the transcriptional hubs. We thank Arif Ul Maula Khan and the Center for Bioimage Analysis (CBA) for their help developing the automated image analysis, and Sarah Kasper from the Data Science Center at EMBL for their suggestions regarding the epistasis analysis. We thank Aliaksandr Halavatyi and Manuel Gunkel, and the Advanced Light Microscopy Facility (ALMF) at EMBL for their help with the imaging. Images of adult Drosophila species used in Figure S2g and i are courtesy of the Obbard Lab (https://obbard.bio.ed.ac.uk/photos.html), licensed under CC BY-NC 4.0. Lastly, we want to thank all the present and past members of the Crocker lab for their input and advice.

