## Supplementary Figures for "Interallelic cis-regulatory dominance promotes robustness and evolutionary innovation"

**The PDF file includes:**
Figs. S1 to S6

**Other Supplementary Materials for this manuscript include the following:**
Suplementary File 1.
Supplementary Table 1.
Supplementary Table 2.

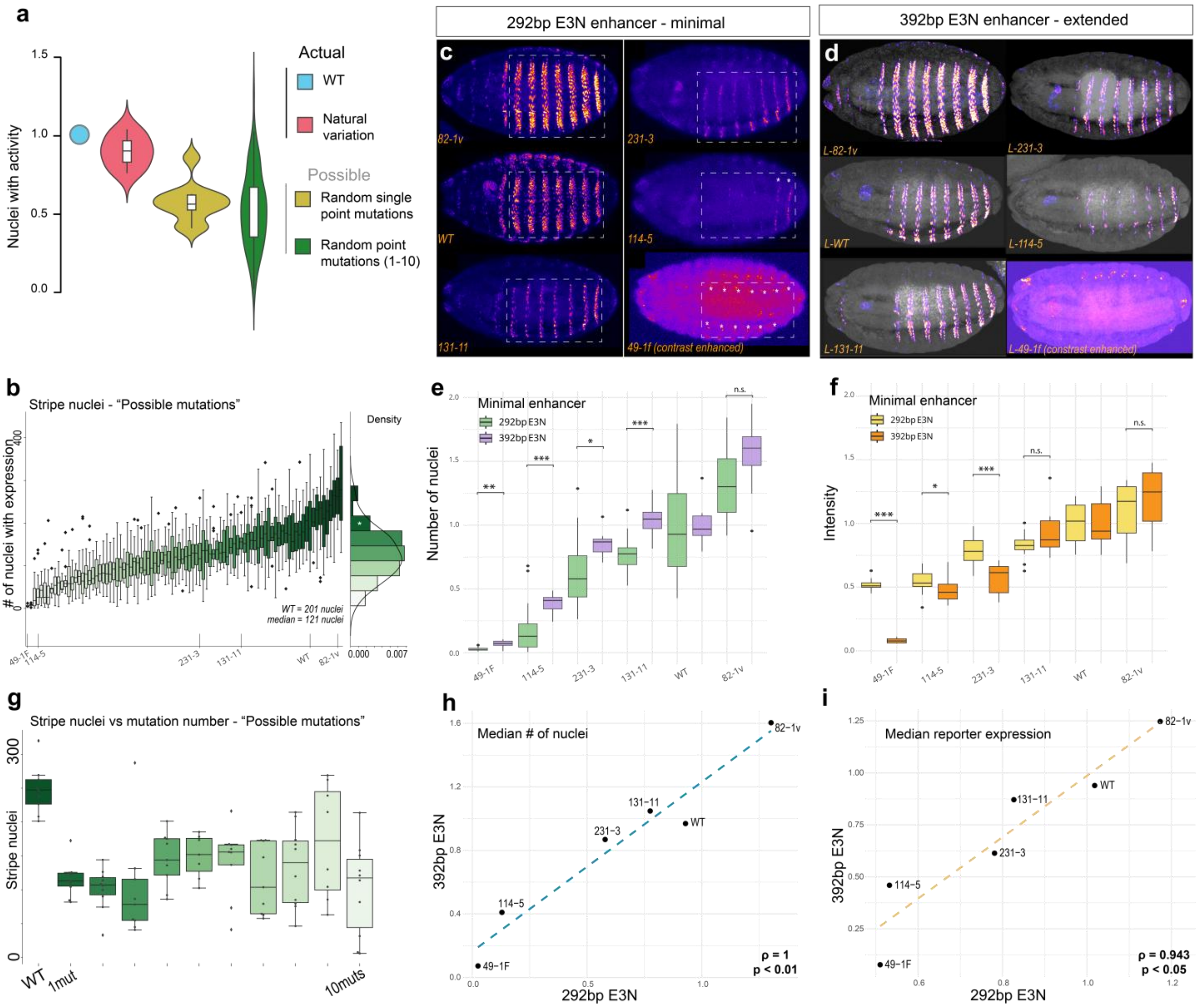


**Supplementary Figure 1. Phenotypic effects of mutations in the minimal *E3N* enhancer are not substantially buffered by the sequences directly flanking it in the native locus.**

(**a**) A comparison of the effect of the natural variation, random single point mutations, and random mutations ranging from 1 to 10 mutations in the *E3N* enhancer on the number of nuclei with *lacZ* expression. Each datapoint in the boxplot represents the average of a genotype.

(**b**) Boxplot showing the number of nuclei with expression of the *lacZ* reporter for the 91 original mutant variants in the 292bp *E3N-lacZ* construct. On the x-axis the selected mutant variants are highlighted.

(**c**) Representative images of embryos from the original mutant variants in the 292bp *E3N-lacZ* construct.

(**d**) Representative images of embryos from the extended ("L") mutant variants in the 392bp *E3N-lacZ* construct.

(**e**) Boxplot showing the number of nuclei with expression of the *lacZ* reporter of different mutant variants of *E3N* in the normal 292bp versus the extended 392bp *E3N-lacZ* construct, normalized to their respective wildtypes. Adjusted p-values (FDR) for the different mutant variants: 49-1F, p = 0.003, 114-5, p = 4.129e-4, 231-3 = 0.011, 131-11, p = 4.129e-4, 82-1v, p = 0.093.

(**f**) Boxplot showing the mean intensity of the *lacZ* reporter of different mutant variants of *E3N* in the minimal 292bp versus the extended 392bp *E3N-lacZ* construct, normalized to their respective wildtypes. Adjusted p-values (FDR) for the different mutant variants: 49-1F, p = 2.706e-07, 114-5, p = 0.044, 231-3 = 3.072e-05, 131-11, p = 0.158, 82-1v, p = 0.424.

(**g**) Boxplot showing the effect of the number of mutations within the 91 original mutant variants in the 292bp *E3N-lacZ* construct on the number of nuclei with expression of the *lacZ* reporter.

(**h**) Correlation of median number of nuclei with *lacZ* reporter expression for every mutant variant in the minimal (x-axis) or extended context (y-axis). The dashed line represents a linear fit (Spearman's ρ = 1, p = 0.003).

(**i**) Correlation of median β-gal intensity for every mutant variant in the minimal (x-axis) or extended genetic context (y-axis). The dashed line represents a linear fit (Spearman's ρ = 0.943, p = 0.017).

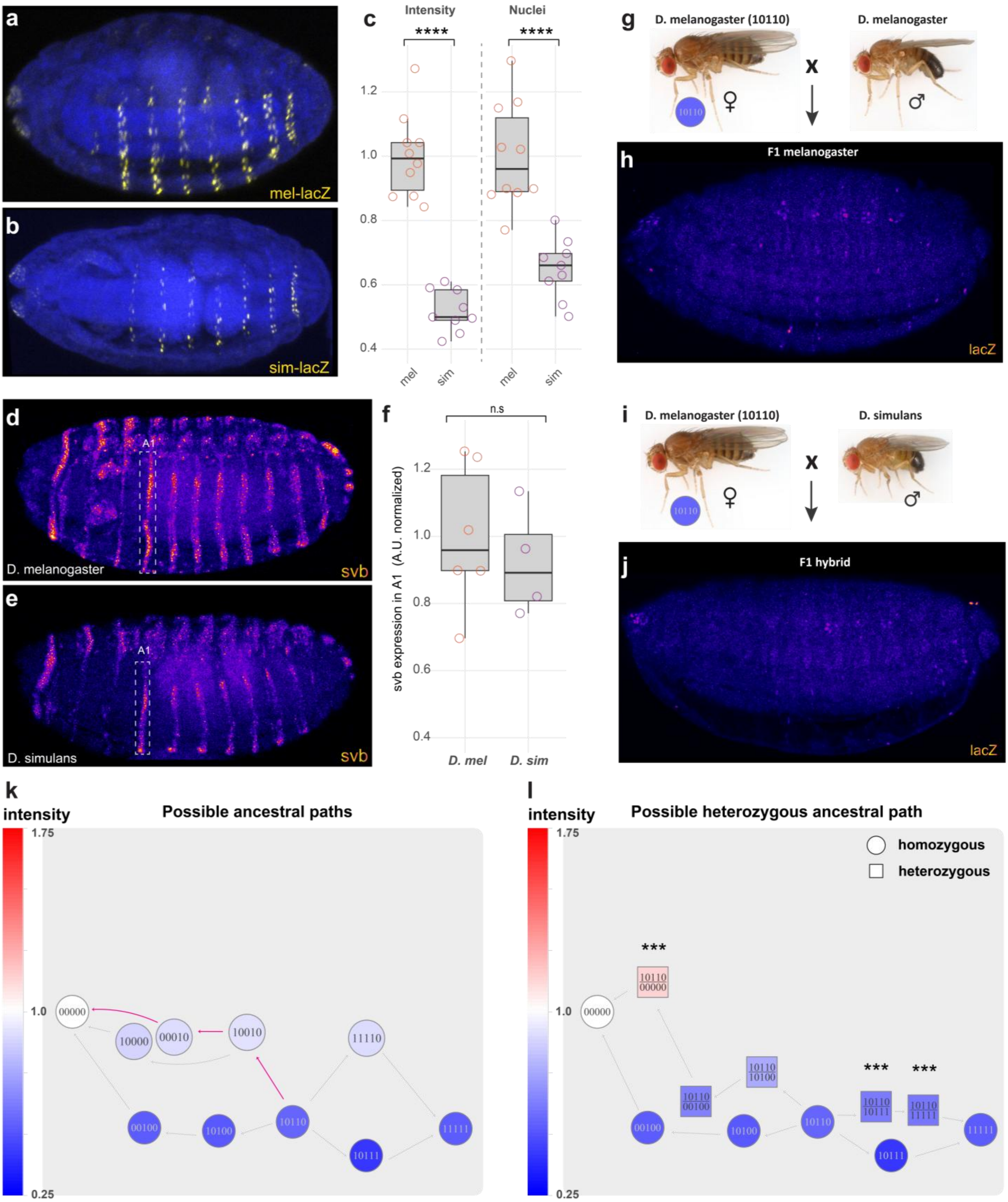


**Supplementary Figure 2. No evidence for divergence in the trans-environment of *E3N* between *D. melanogaster* and *D. simulans*.**

(**a**) *lacZ* expression pattern of a representative *mel-lacZ* embryo from the combinatorial *E3N* library.
(**b**) *lacZ* expression pattern of a representative *sim-lacZ* embryo from the combinatorial *E3N* library.
(**c**) Boxplot showing the difference in nuclear β-gal intensity ($p = 1.43 \times 10^{-8}$) and the number of β-gal positive nuclei ($p = 3.03 \times 10^{-5}$) of the lines shown in (a) and (b). Pairwise comparisons were performed using Tukey's HSD test (confidence level 0.95).
(**d**) HCR-RNA FISH staining of a stage 15 *D. melanogaster* embryo with a probe for the first exon of *svb* (n=6).
(**e**) HCR-RNA FISH staining of a stage 15 *D. simulans* embryo with a probe for the first exon of *svb* (n=4).

(**f**) Quantification of *svb* expression in the A1 region of embryos from D (n=6) and E (n=4), normalized to *D. melanogaster*, showing no significant difference between *svb* expression between the two species (p = 0.558). Pairwise comparisons were performed using Tukey's HSD test (confidence level 0.95).

(**g**) Overview of the cross performed between a *D. melanogaster* female homozygous for the *10110-lacZ* construct and a *D. melanogaster* male without any reporter construct inserted. Images of the *D. melanogaster* adult flies adapted from the Obbard Lab, used under a Creative Commons CC BY-NC 4.0 license.

(**h**) *lacZ* expression pattern of a representative embryo from a cross described in (g). F1 offspring (as pictured) carry a single copy of the ancestral 10110 version of the *E3N-lacZ* construct.

(**i**) Overview of the cross performed between a *D. melanogaster* female homozygous for the *10110-lacZ* construct and a *D. simulans* male without any reporter construct inserted. Images of the *D. melanogaster* and *simulans* adult flies adapted from the Obbard Lab, used under a Creative Commons CC BY-NC 4.0 license.

(**j**) *lacZ* expression pattern of a representative embryo from a cross described in (i). F1 hybrid *D. melanogaster-simulans* offspring (as pictured) carry a single copy of the ancestral *10110* version of the *E3N-lacZ* construct.

(**k**) Possible ancestral paths from *10110* to modern-day *D. simulans* and *D. melanogaster* based on the alignment to the modern-day *D. erecta E3N* enhancer*.* Arrows in pink represent a more likely path, as the *00010* variant is found in modern-day *D. melanogaster* populations as a SNP (see **Table S1**).

(**l**) A comparison of one of the evolutionary trajectories from the suspected ancestral form of *E3N* (*10110*) to modern-day mel and sim when every step is homozygous versus heterozygous, where one allele remains ancestral (*10110*) up until the last step towards *mel* and *sim*. The nodes in these paths are positioned on a y-axis relative to their normalized average intensity of the nuclei with enhancer activity. Linear models were fit independently for each cross that produced heterozygotes. Dominance deviations, defined as a significant p-value for βd, are marked with asterisks (* for p < 0.05, ** for p < 0.01, *** for p < 0.001). All non-significant heterozygote nodes are assumed to fit an additive model (βa). See **Table S2** for a full description of the linear models.

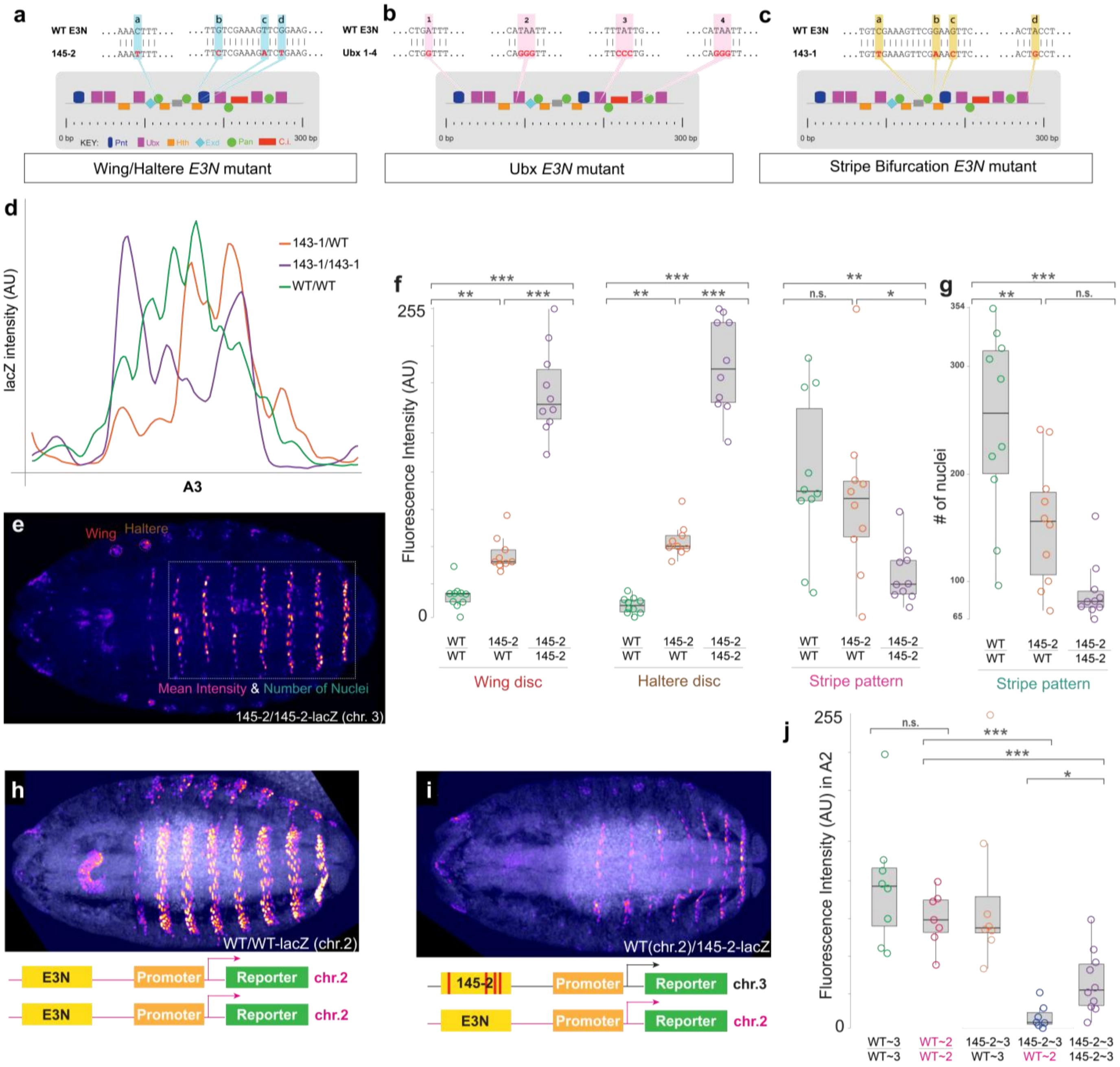


**Supplementary Figure 3. Interallelic buffering capacity is impaired when alleles are placed on different chromosomes.**
(**a**) Schematic of the mapped binding sites of the *E3N* enhancer. The nucleotides mutated in the *145-2* construct are marked in blue.
(**b**) Schematic of the mapped binding sites of the *E3N* enhancer. The nucleotides mutated in the Ubx sites in the *Ubx1-4* construct are marked in pink.
(**c**) Schematic of the mapped binding sites of the *E3N* enhancer. The nucleotides mutated in the *143-1* construct are marked in yellow.
(**d**) Line graph showing the β-gal reporter intensity profiles of the representative embryos depicted in Figure 3 (g-i) measured in the white box (stripe A3).
(**e**) A representative stage 15 embryo of the homozygous *145-2 E3N* variant, stained with a β-gal antibody against the *lacZ* reporter protein β-gal. Different zones measured in (f) and (g) depicted in red (primordium wing disc), brown (primordium haltere disc), magenta and cyan (mean β-gal intensity and number of nuclei with *lacZ* expression from A2 to A8).
(**f**) Boxplots of the mean fluorescence intensity of the β-gal reporter within the wing disc, haltere disc and ventral zone (stripe A2 to A8) of different embryos of the *145-2 E3N* variant at stage 15. Pairwise comparisons were performed using Tukey's HSD test (confidence level 0.95). For the Wing disc boxplot, all group comparisons were significant: WT/WT vs. 145-2/WT, p = 0.010; WT/WT vs. 145-2/145-2, p = 4.87 × $10^{-14}$; 145-2/WT vs. 145-2/145-2, p = 7.04 × $10^{-12}$. For the Haltere disc boxplot, all group comparisons were significant: WT/WT vs. 145-2/WT, p = 1.81 × $10^{-4}$; WT/WT vs. 145-2/145-2, p = 3.10 × $10^{-14}$; 145-2/WT vs. 145-2/145-2, p = 2.80 × $10^{-12}$. For the Stripe pattern boxplot, two out of three comparisons were significant: WT/WT vs. 145-2/WT, p = 0.738; WT/WT vs. 145-2/145-2, p = 8.81 × $10^{-3}$; 145-2/WT vs. 145-2/145-2, p = 4.95 × $10^{-2}$.
(**g**) Boxplot of the number of nuclei with *lacZ* reporter expression in the ventral zone (stripe A2 to A8) of different embryos of the

*145-2 E3N* variant at stage 15. Pairwise comparisons were performed using Tukey's HSD test (confidence level 0.95). Two out of three comparisons were significant: WT/WT vs. 145-2/WT, p = 8.25 × $10^{-3}$; WT/WT vs. 145-2/145-2, p = 2.25 × $10^{-5}$; 145-2/WT vs. 145-2/145-2, p = 8.25 × $10^{-3}$.
(**h**) Representative stage 15 embryo of a homozygous wildtype *E3N-lacZ* line, integrated at *attP40* (chr.2). Embryos are stained with a β-gal antibody against the *lacZ* reporter protein β-gal.
(**i**) Representative stage 15 embryo of a heterozygous line, resulting from a cross of the wildtype *E3N-lacZ* line, integrated at *attP40* (chr.2) with ectopic *E3N* mutant *145-2*, integrated at *attP2* (chr.3L). Embryos are stained with a β-gal antibody against the *lacZ* reporter gene.
(**j**) Boxplot showing the fluorescence intensity in the A2 stripe region of the β-gal reporter of different crosses with *E3N-lacZ* integrated at either *attP2* (chr.3L) or *attP40* (chr.2). Pairwise comparisons were performed using Tukey's HSD test (confidence level 0.95): WT~3/WT~3 vs. WT~2/WT~2 p = 0.101; WT~2/WT~2 vs. 145-2~3/WT~2, p = 2.04 × $10^{-6}$; WT~2/WT~2 vs. 145-2~3/145-2~3, p = 2.22 × $10^{-4}$; 145-2~3/WT~2 vs. 145-2~3/145-2~3, p = 3.67 × $10^{-2}$.

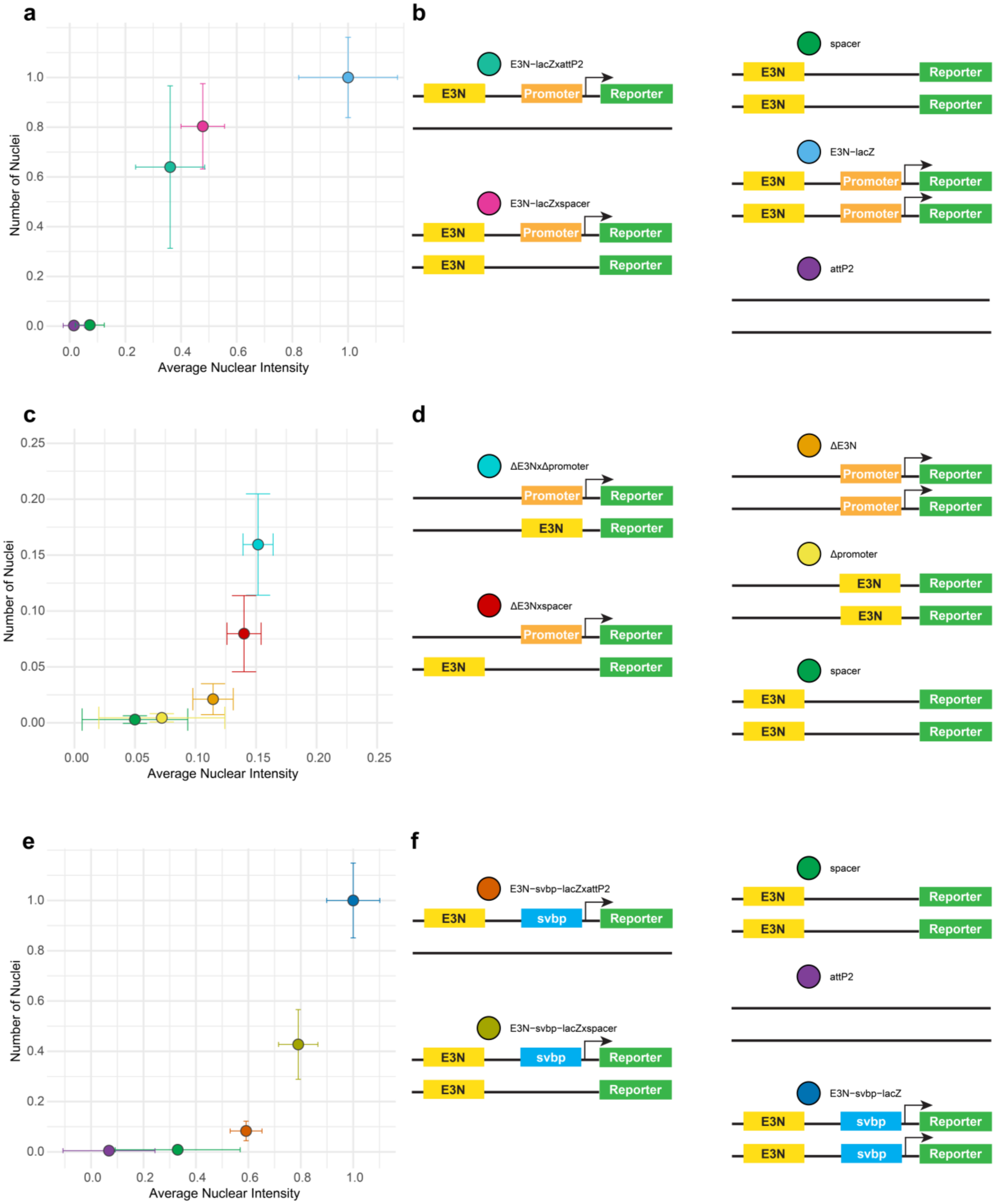


**Supplementary Figure 4. Interallelic compensation persists when the *hsp70* promoter is hemizygously removed, or switched with the native *svb* promoter.**
(**a**) Scatterplot, where points represent the mean of normalized nuclear β-gal intensity and number of nuclei with β-gal signal for different genotypes, with error bars representing 95% confidence intervals of the mean. All genotypes are normalized to the *E3N-lacZ* genotype (blue). Different colors represent the genotypes pictured in (b).
(**b**) Schematics of the different genotypes represented in the scatterplot in (a), where the insertion at the *attP2* site on the third chromosome in *D. melanogaster* is shown.

(**c**) Scatterplot, where points represent the mean of normalized nuclear β-gal intensity and number of nuclei with β-gal signal for different genotypes, with error bars representing 95% confidence intervals of the mean. All genotypes are normalized to the *E3N-lacZ* genotype (blue). Different colors represent the genotypes pictured in (d).
(**d**) Schematics of the different genotypes represented in the scatterplot in (c), where the insertion at the *attP2* site on the third chromosome in *D. melanogaster* is shown.
(**e**) Scatterplot, where points represent the mean of normalized nuclear β-gal intensity and number of nuclei with β-gal signal for different genotypes, with error bars representing 95% confidence intervals of the mean. All genotypes are normalized to the *E3N-svbp-lacZ* genotype (dark blue). Different colors represent the genotypes pictured in (f).
(**f**) Schematics of the different genotypes represented in the scatterplot in (e), where the insertion at the *attP2* site on the third chromosome in *D. melanogaster* is shown.

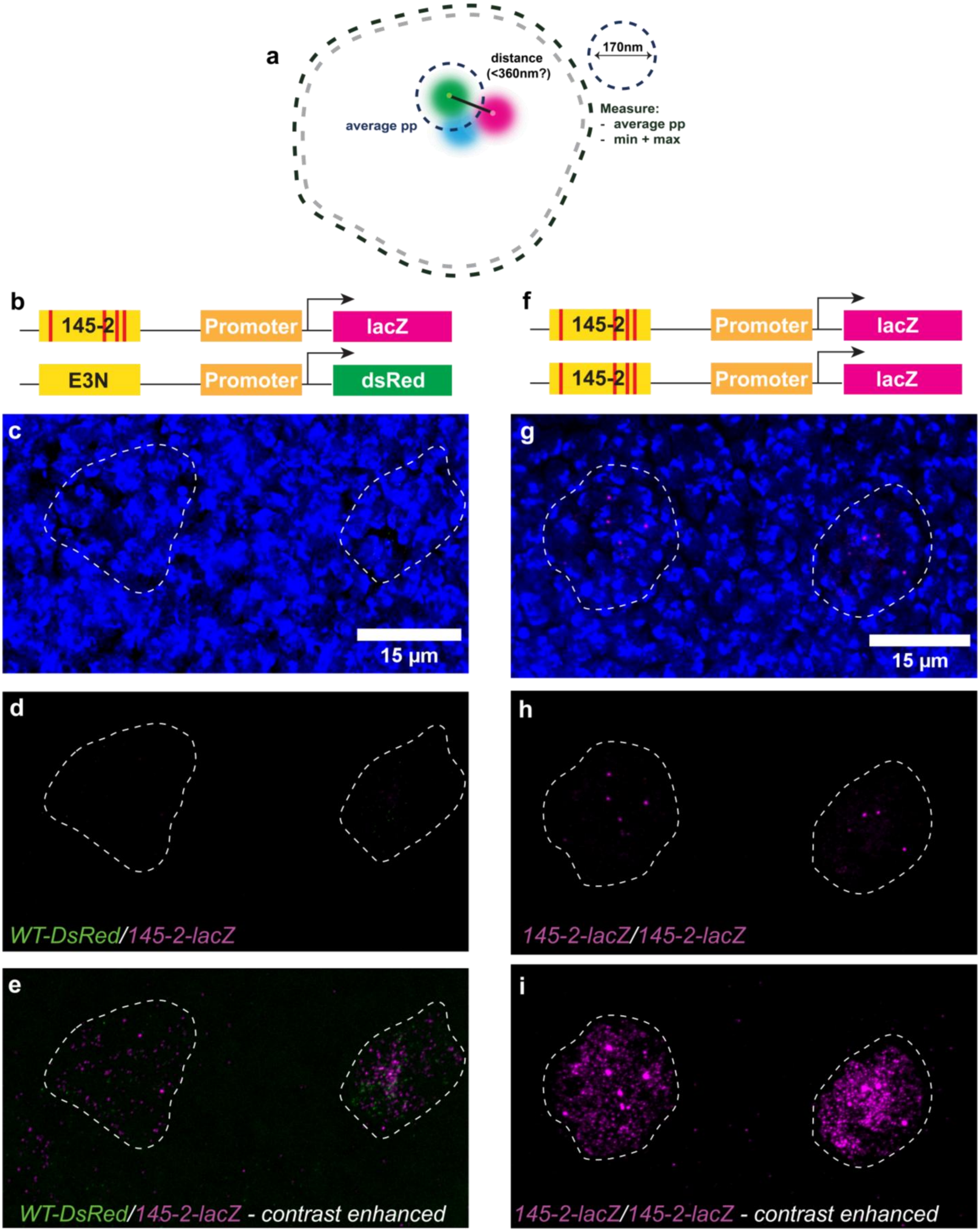


**Supplementary Figure 5. Differential transcription in the imaginal discs of homozygous and heterozygous 145-2 embryos.**
(**a**) Graphical depiction of the different measurements performed on the two active reporter transcription sites (green and magenta) and the Ubx signal (light blue) within a nucleus. pp = per pixel. See **Methods** for a full description.

(**b**) Schematic of the heterozygous *145-2-lacZ/E3N-dsRed* genotype shown in (c-e). Distinct reporter genes allow allele-specific visualization of transcriptional activity.

(**c**) Close-up of the wing (left) and haltere (right) discs of a stage 15 embryo corresponding to (b), indicated by dotted circles in **Fig. 4g**. Nuclei are labeled with DAPI (blue).

(**d**) Same close-up as in (c), shown without the DAPI channel. Magenta represents *lacZ* signal from HCR RNA-FISH staining; spots correspond to nascent RNA at active *lacZ* transcription sites.

(**e**) Same as (d), with contrast digitally enhanced to reveal both nascent transcripts at transcription sites and mature *lacZ* RNA molecules.

(**f**) Schematic of the homozygous *145-2-lacZ* genotype shown in (g-i).

(**g**) Close-up of the wing (left) and haltere (right) discs of a stage 15 embryo corresponding to (f). Nuclei are labeled with DAPI (blue)

(**h**) Same close-up as in (g), shown without the DAPI channel. Green represents *dsRed* signal, and magenta represents *lacZ* signal from HCR RNA-FISH staining.

(**i**) Same as (h), with contrast digitally enhanced to reveal mature *lacZ* and *dsRed* RNA molecules.

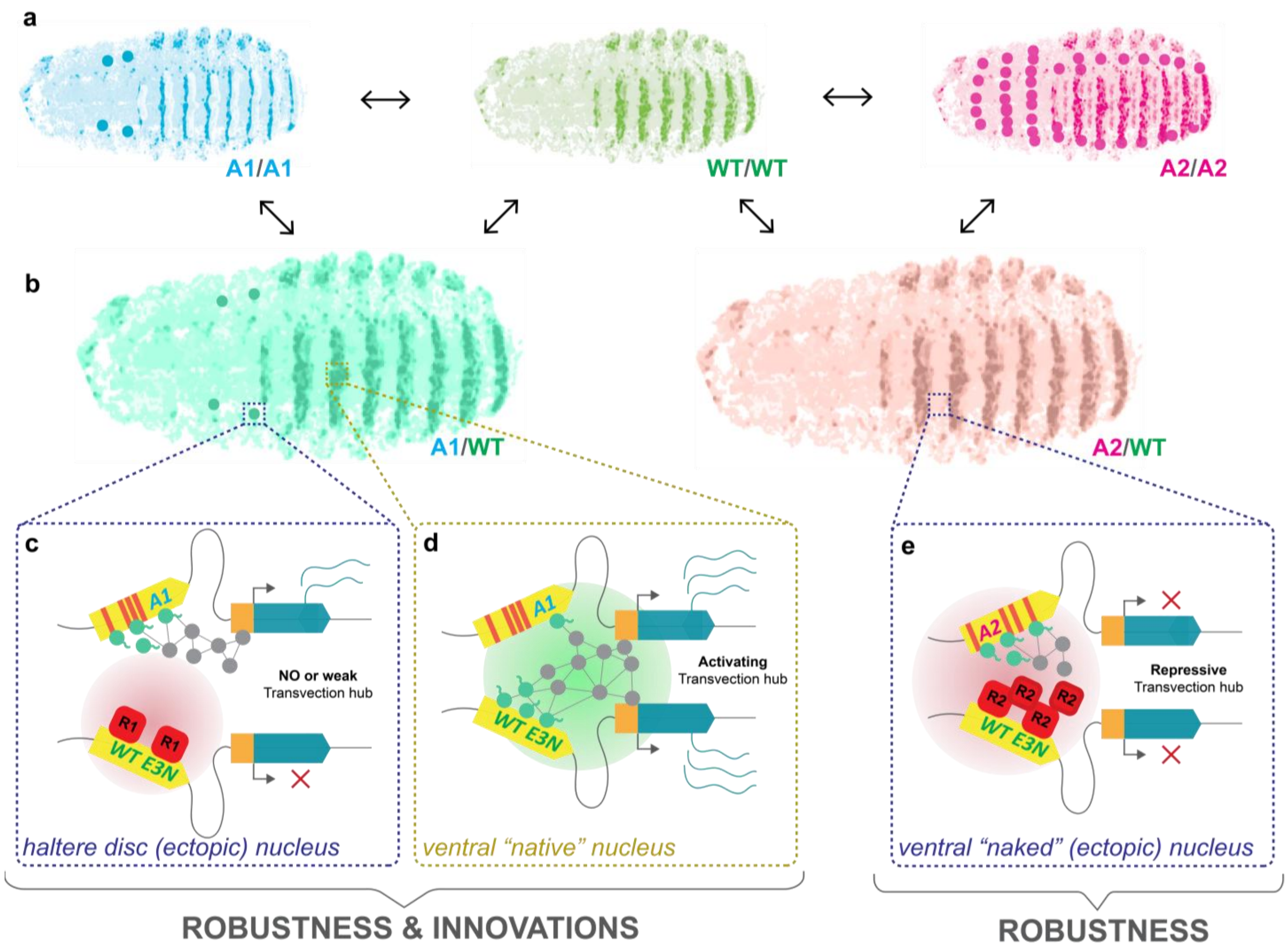


**Supplementary Figure 6. Conceptual model of how mosaic regulatory dominance could lead to robustness and innovations in *Drosophila*.**

(**a**) Graphical depiction of the homozygous effects of allele 1 (A1) and allele 2 (A2) on a gene expression pattern in a late-stage *Drosophila* embryo.

(**b**) Graphical depiction of the heterozygous effects of A1 or A2 on a gene expression patten in a late-stage *Drosophila* embryo, where different cell types exhibit different degrees of dominance in gene expression.

(**c**) Graphical depiction of a close-up of the locus in the haltere of the left embryo described in (b). Here, repressors binding at the WT enhancer lead to no, or a weak repressive transvection hub. This regulatory architecture could explain a mild underdominant effect on gene expression in this cell type.

(**d**) Graphical depiction of a close-up of the locus in the ventral ectodermal zone of the left embryo described in (b). Here, mutations lead to a lack of activation from A1, but this is compensated by a transvection hub. This regulatory architecture is consistent with dominance of the wildtype allele in this cell type.

(**e**) Graphical depiction of a close-up of the locus in the "naked" ventral ectodermal zone of the right embryo described in (b). Here, repressors binding at the WT enhancer lead to a strong repressive transvection hub. This regulatory architecture could explain the dominant repressive effect on gene expression in these cells relative to the A2/A2 embryo.
